# Asymmetric cell division-specific phosphorylation of PAR-3 regulates neuroblasts polarisation and sensory organ formation in *Drosophila*

**DOI:** 10.1101/2023.07.26.550680

**Authors:** Nicolas Loyer, Elizabeth K. J. Hogg, Hayley Shaw, David H. Murray, Greg M. Findlay, Jens Januschke

## Abstract

The generation of distinct cell fates during development depends on asymmetric cell division of progenitor cells. In the central and peripheral nervous system of *Drosophila,* progenitor cells respectively called neuroblasts or sensory organ precursors use PAR polarity during mitosis to control cell fate determination in their daughter cells. How polarity and the cell cycle are coupled, and how the cell cycle machinery regulates PAR protein function and cell fate determination is poorly understood. Here, we generate an analog sensitive allele of CDK1 and reveal that its partial inhibition weakens but does not abolish apical polarity in embryonic and larval neuroblasts, and leads to defects in polarisation of fate determinants. We describe a novel *in vivo* phosphorylation of Bazooka, the *Drosophila* homolog of PAR-3, on Serine180, a consensus CDK phosphorylation site. Remarkably, phosphorylation of Serine180 occurs in asymmetrically dividing neuroblasts and sensory organ precursors, and not in their symmetrically dividing neighbours. We further show that Serine180 phosphomutants disrupt the timing of basal polarisation in neuroblasts and sensory organ formation in sensory organ precursors. Finally, we show that CDK1 can phosphorylate human PARD3 *in vitro,* suggestive of a conserved kinase-substrate relationship between CDK1 and PAR-3.

## Introduction

The coordination of cellular events in time and space during cell division is important for development. This spatiotemporal organization is particularly critical for certain types of asymmetric cell division that generate different cell fates in a single division. Cell fate difference in the resulting daughter cells can stem, for instance, from the asymmetric subcellular localisation of fate determinants in the dividing cell. This asymmetry results in the daughter cell receiving distinct sets of molecular information. In such divisions, the subcellular localisation and segregation of molecules involved in fate decisions needs to be timed along the cell cycle for proper establishment of cells fates of the resulting daughter cells. The molecular mechanisms underpinning this coordination are not fully resolved.

Studies in worms and in flies have uncovered key aspects into the coupling of the cell cycle and cell fate and suggest that the cell cycle dependent regulation of cell polarity is important in this context (Noatynska et al., 2013, Fichelson et al., 2005). In many asymmetric dividing cells the conserved PAR polarity complex, composed of PAR-3, PAR-6 and atypical protein kinase C (aPKC) (Goldstein and Macara, 2007, Goehring, 2014), polarises prior to division. The PAR polarity complex exhibits cell-cycle dependent localisation in the*C. elegans* zygote (Kemphues, 2000) and many stem and precursor cells, such as neural precursors in the chick (Das and Storey, 2012) and fish (Alexandre et al., 2010), muscle stem cells in the mouse (Dumont et al., 2015), and stem and precursor cells of the developing central and peripheral nervous system of *Drosophila* (Wodarz et al., 1999, Schober et al., 1999, Bellaiche et al., 2001). The mechanisms controlling the subcellular localisation and function of the PAR complex in cycling cells are poorly understood.

In the *C.elegans* zygote, the coordination of cell polarity by the cell cycle machinery is critical for a successful asymmetric division establishing the precursor of the germ line and that of somatic fates in a single division (Rose and Kemphues, 1998). Cell cycle kinases like Plk1 and Aurora A (Kim and Griffin, 2020, Reich et al., 2019, Schumacher et al., 1998) regulate polarity in the zygote required for the establishment of PAR anterior-posterior polarisation. In flies, the homologs of Plk1 and Aurora A and the master regulator of cell cycle transitions, CDK1 (Nurse, 1997), have been implicated in linking cell cycle and asymmetric division in the central and peripheral nervous system (Tio et al., 2001, Wirtz-Peitz et al., 2008, Lee et al., 2006, Wang et al., 2006, Wang et al., 2007).

*Drosophila* neural stem cells called neuroblasts are a well-studied model for asymmetric cell division (Sunchu and Cabernard, 2020, Gallaud et al., 2017). Neuroblasts divide asymmetrically generating a self-renewed neuroblast and a differentiating daughter cell. This outcome relies on a series of events taking place during mitosis. In prophase, PAR proteins (PAR-6, aPKC and PAR-3, known as Baz in *Drosophila*) accumulate to a cortical domain, defining the apical pole (Rolls et al., 2003, Wodarz et al., 1999, Wodarz et al., 2000, Petronczki and Knoblich, 2001). The earliest signs of neuroblast polarisation in mitosis include the recruitment of Baz from the cytoplasm to a broad apical area at the onset of prophase (Hannaford et al., 2018). At nuclear envelope breakdown (NEB), basal-to-apical actin-driven cortical flows then lead to the coalescence of Baz into continuous a continuous cluster that forms a bright apical crescent (Oon and Prehoda, 2019). The signals that drive Baz coalescence and its functional significance in neuroblasts are unknown.

Following nuclear envelope break down, the PAR complex excludes cell fate determinants such as Miranda, Pon and Numb from the apical pole *via* aPKC activity, leading to their localisation at the opposite basal pole. At metaphase, the mitotic spindle aligns with the apical basal polarity axis, leading to the asymmetric segregation of fate determinants into the differentiating daughter cell (Gillies and Cabernard, 2011). Therefore, cell polarity and the cell cycle are tightly coordinated in neuroblasts, with apical polarity assembled and disassembled at each cell cycle. Baz is heavily regulated by phosphorylation by different kinases including PAR-1 and aPKC in flies (Benton and St Johnston, 2003, Morais-de-Sá et al., 2010) and additionally by Aurora A, Plk1 and other kinases in other systems (Dickinson et al., 2017, Khazaei and Püschel, 2009). Thus, cell cycle-dependent kinases, indispensable for the spatio-temporal regulation of cell divisions, are promising candidates for coordinating polarity with the cell cycle through direct phosphorylation of Baz.

Some polarity proteins are phosphorylated by cell cycle dependent kinases during asymmetric neuroblast division: Aurora A phosphorylates PAR-6 (Wirtz-Peitz et al., 2008) and Polo, the fly homolog of Plk1, phosphorylates Pon (Wang et al., 2007). Importantly, although these phosphorylation events control basal polarity, apical Baz polarity remains largely unaffected in *aurora A* and *polo* mutants (Lee et al., 2006, Wang et al., 2006, Wang et al., 2007). Interestingly, Baz asymmetric localisation was reported to be lost in embryonic neuroblasts expressing a dominant negative form of CDK1 (Tio et al., 2001). Thereby, direct phosphorylation of Baz by CDK1 may be the temporal trigger driving Baz polarisation in mitotic neuroblasts.

Like in neuroblasts, Baz polarity is coupled to the cell cycle in asymmetrically dividing sensory organ precursors (SOPs) in the peripheral nervous system of the fly (Bellaiche et al., 2001). In this system, SOP cells are first specified through a Notch-dependent lateral inhibition mechanism (Simpson, 1990), and then undergo a series of asymmetric cell divisions generating five cells. At the end of each division, the asymmetric segregation of Notch regulators results in the differential activation of the Notch pathway in each daughter cell, assigning them different identities (Schweisguth, 2015). Downregulation of CDK1 activity in SOPs delays mitosis and causes mother-to-daughter cell fate transformations occurring without divisions, affecting polarity orientation when cells eventually divide, and ultimately leads to loss of external organs (Fichelson and Gho, 2004). Interestingly, Cyclin A, a CDK1 activator, was recently found to colocalise with Baz at the posterior cortex of mitotic SOPs (Darnat et al., 2022), consistent with the possibility that CDK1 may directly phosphorylate Baz.

Here, we used live cell imaging and chemical genetics to study the cell cycle and cell polarity coordination in *Drosophila* neuroblasts and SOPs. We generated an analog-sensitive allele of CDK1 allowing us to tune CDK1 activity. Partial CDK1 inhibition in larval neuroblasts affects Baz polarity by preventing apical crescent coalescence at NEB. We identified a consensus CDK phosphorylation on the Baz sequence (Baz-S180), against which we developed a phospho-specific antibody. We found Baz-S180 to be phosphorylated specifically during the asymmetric division of neuroblasts and sensory organ precursors but not during the symmetric cell divisions of neighbouring cells in either context. We further analysed Baz-S180 phosphomutants and observed that, despite not being necessary for Baz localisation, phosphorylation of Baz-S180 controls the timing of basal polarisation in neuroblasts and the specification of sensory organs. Finally, we report that human PARD3 is a substrate for CDK1/Cyclin B *in vitro*.

## Results

### Generation and *in vivo* analysis of an analog-sensitive allele of CDK1

To investigate the role of CDK1 in regulating Baz localization during the neuroblast cell cycle we generated an analog sensitive allele (Lopez et al., 2014) of *cdk1*, *cdk1^F80A^*, henceforth called *cdk1^as2^*. To test whether CDK1 activity can be acutely and specifically inhibited in these mutants, we exposed control and *cdk1^as2^* larval brains to the ATP analog 1-NA-PP1, using cell cycle arrest as an indicator (Nurse, 1997). As we previously showed that 10 µM of 1-NA-PP1 acutely blocks an analog sensitive version of aPKC (Hannaford et al., 2019), we used 10 µM as a starting concentration. While control neuroblasts continued proliferating after exposure to 1-NA-PP1, *cdk1^as2^* neuroblasts stopped dividing (**Fig 1A,A’**). Interestingly, upon 1-NA-PP1 addition, early prophase *cdk1^as2^* neuroblasts lost their apical Baz crescent and did not further proceed through mitosis (**Fig 1B**), apparently reverting back to interphase, a reported effect of CDK1 inhibition (Potapova et al., 2011). Another expected effect of CDK1 inhibition is exit from mitosis in metaphase-arrested cells (mitotic exit) (Potapova et al., 2006). We exposed metaphase-arrested neuroblasts to 1-NA-PP1, which resulted in mitotic slippage within 10 minutes in the majority of *cdk1^as2^* neuroblasts, but not in controls (**Fig 1C,C’**). Interestingly, in neuroblasts that exited mitosis, Baz asymmetry was lost in a manner reminiscent of localization changes normally occurring with the onset of anaphases of untreated neuroblasts. Like in anaphase, Baz crescents spread along the lateral cortex while the cell elongated along the apico-basal axis, after which Baz eventually dissipated into the cytoplasm (**Fig 1D**). We conclude that CDK1 activity is acutely and specifically inhibited in *cdk1^as2^*mutant flies by 10 µM of 1-NA-PP1.

**Figure 1:**
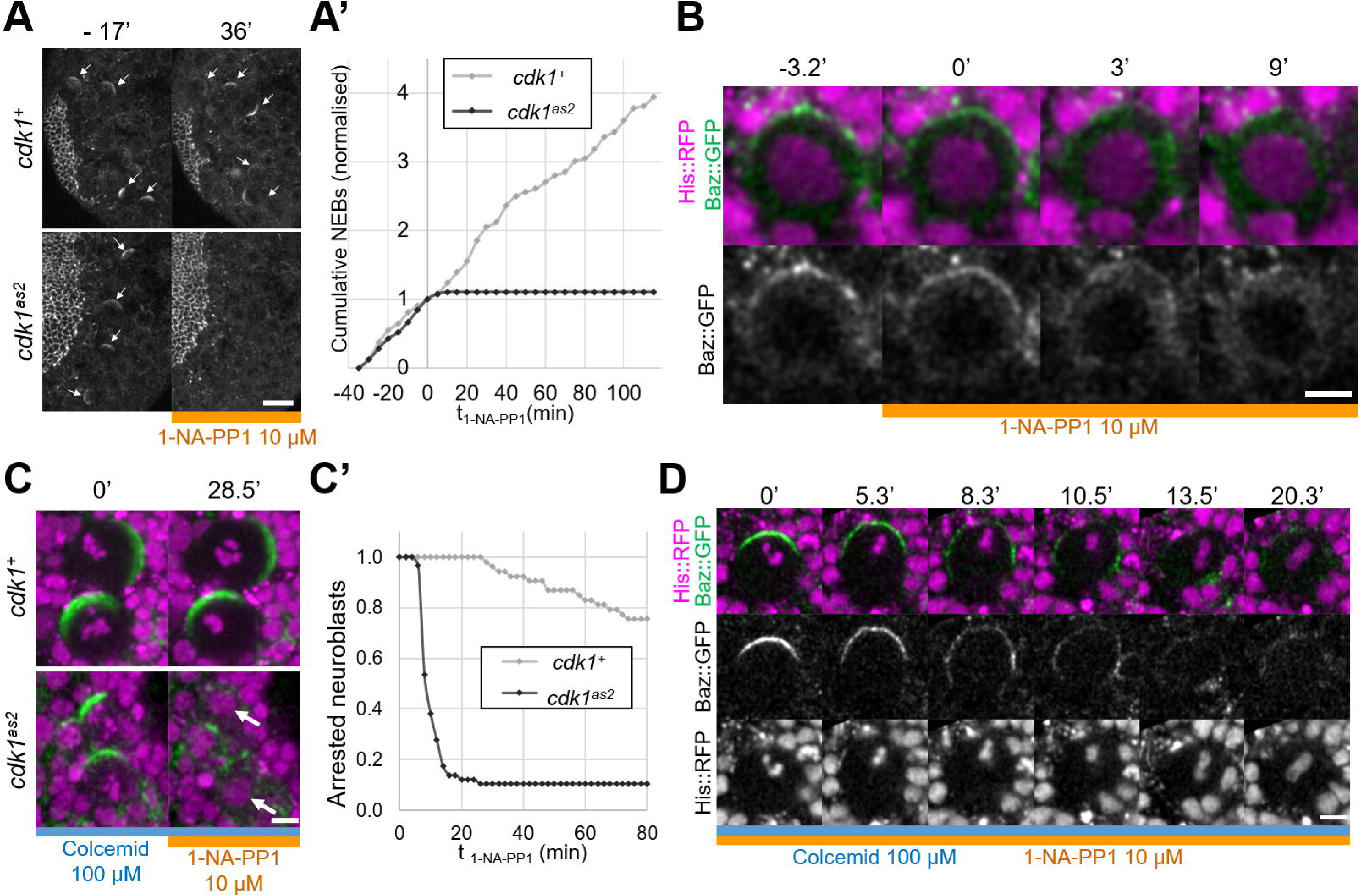
Full inhibition of analog-sensitive CDK1. **A**) Live larval brains expressing Baz::GFP, before and after addition of 1-NA-PP1 10 µM at t0. Arrows point to mitotic neuroblasts. Scale bar: 20 µm. **A’**) Cumulative sum chart of NEB events, normalized to the total number by the time of 1-NA-PP1 addition (t0). N=324 divisions in 4 brains (*cdk1^+^*) and 75 divisions in 5 brains (*cdk1^as2^*), 3 experiments. **B**) Loss of Baz::GFP cortical localization in a cycling *cdk1^as2^* neuroblast at prophase, after addition of 1-NA-PP1 10 µM at t0. Observed in 12 cases across 8 experiments. Scale bar: 5 µm. **C**) Live metaphase-arrested neuroblasts expressing Baz::GFP (green) and His::RFP (magenta), before and after addition of 1-NA-PP1 10 µM at t0. Scale bar: 5 µm. **C’**) Number of metaphase-arrested neuroblasts, normalised to their number at the time of 1-NA-PP1 10 µM addition (t0). N=53 neuroblasts in 5 brains (*cdk1^+^*) and 58 neuroblasts in 5 brains (*cdk1^as2^*), 2 experiments.**D**) Mitotic slippage of a metaphase-arrested *cdk1^as2^* neuroblast after addition of 1-NA-PP1 10 µM at t0. Scale bar: 5 µm.

### Partial inhibition of CDK1 prevents coalescence of apical Baz crescents

We next investigated whether acute CDK1 inhibition affects neuroblasts polarity. As CDK1 inhibition with 10 µM 1-NA-PP1 prevents neuroblasts from cycling and causes metaphase-arrested neuroblasts to slip out of mitosis. This outcome prevents the analysis of polarizing or polarized neuroblasts. Therefore, we partially inhibited Cdk1 using lower concentration of 1-NA-PP1. *cdk1^as2^* neuroblasts stopped proliferating when exposed to 1 µM or higher concentrations but continued proliferating at a slower rate in the presence of 0.5 µM (**Fig S2**). Hence, 10 µM 1-NA-PP1 likely fully inhibits CDK1 and 0.5µM partially inhibits CDK1. We compared apical polarity in mitosis of control and *cdk1^as2^* neuroblasts before and after exposure to 0.5 µM 1-NA-PP1. In control cells, Baz polarized in a dynamic manner. First, it was recruited to a broad cortical crescent during prophase, which then coalesced into a smaller, brighter crescent at NEB, as previously described for unperturbed neuroblasts (Hannaford et al., 2018, Oon and Prehoda, 2019). Interestingly, although Baz still polarized upon partial CDK1 inhibition in *cdk1^as2^* neuroblasts, apical Baz crescents appeared fainter and wider than during the previous cell cycle occurring before exposure to 1-NA-PP1 (**Fig 2A, A’**). Thus, Baz crescents fail to coalesce at NEB upon partial CDK1 inhibition. As Baz polarity was previously shown to be completely lost in dominant negative *cdk1* mutant embryonic neuroblasts (Tio et al., 2001), we tested the effect of partial inhibition of CDK1 in larval neuroblasts in a sensitized context. Baz localization is dependent on an oligomerization domain (OD) and a lipid binding domain (LD), which are functionally redundant (Kullmann and Krahn, 2018). We reasoned that deletion of either of these domains might make Baz localization more sensitive to disruptions and lead to complete loss of polarity upon partial inhibition of CDK1. However, Baz deletion mutants for either of these domains still polarized when we partially inhibited CDK1, and only presented an apical coalescence defect (**Fig 2B**). To test whether embryonic neuroblasts are more sensitive to CDK1 inhibition than larval neuroblasts, we exposed embryonic neuroblasts to 0.5 µM 1-NA-PP1. Again, partial inhibition of CDK1 in embryonic neuroblasts resulted in defective coalescence but not in complete loss of polarity (**Fig 2C**). We conclude that high levels of CDK1 activity are needed for full apical polarization of embryonic and larval neuroblasts by driving apical coalescence.

**Figure 2:**
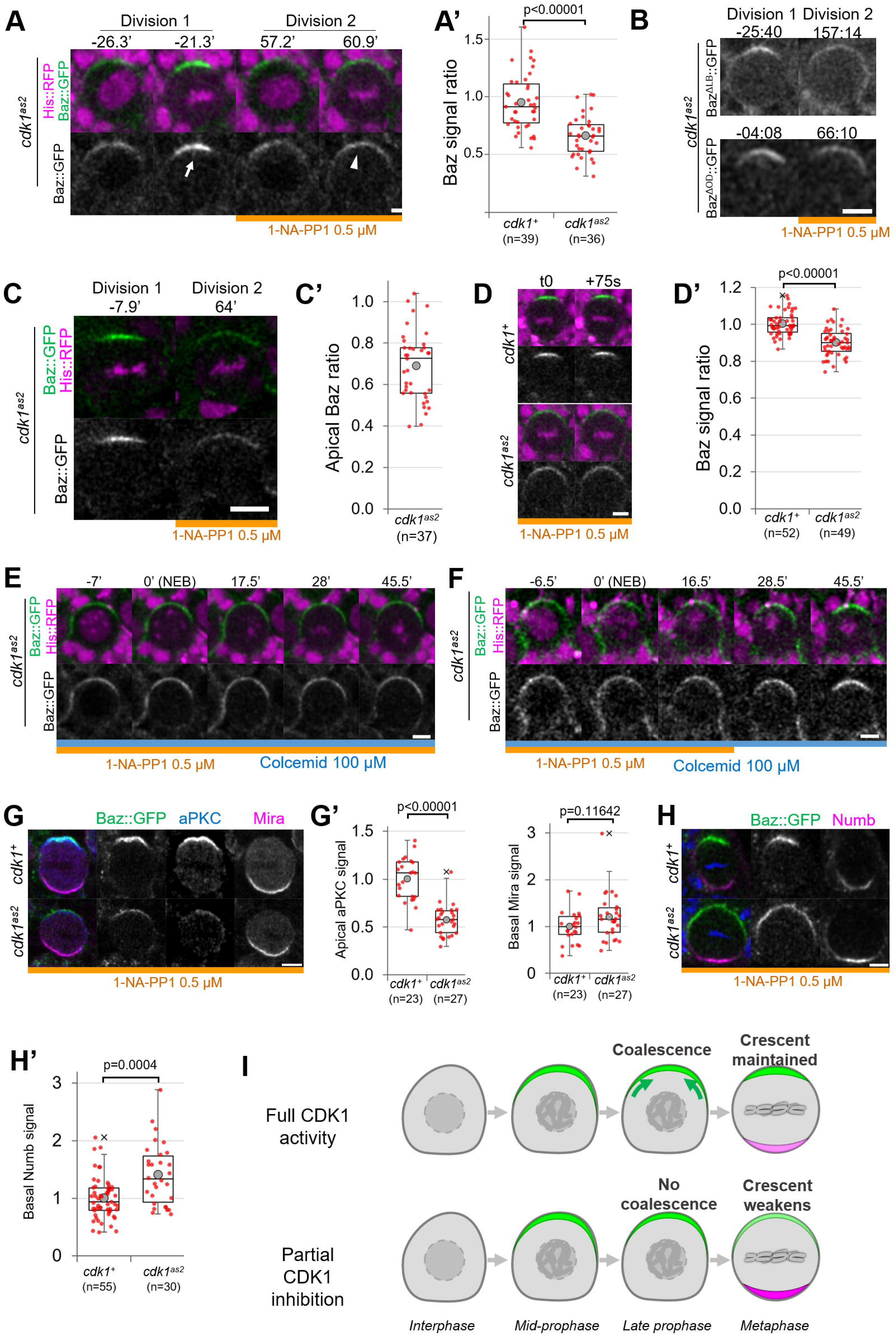
Partial inhibition of analog-sensitive CDK1. **A**) Two consecutive divisions of a live *cdk1^as2^* cycling neuroblast expressing Baz::GFP, before and after addition of 1-NA-PP1 0.5 µM at t0. Arrow: coalesced Baz crescent in metaphase before 1-NA-PP1 addition. Arrowhead: non-coalesced crescent in the next metaphase, in the presence of 1-NA-PP1. Scale bar: 5 µm. **A’**) Ratio between the apical Baz signal at metaphase in the presence of 1-NA-PP1 0.5 µM and the Baz signal during the previous metaphase, in the absence of 1-NA-PP1. In control neuroblasts, 1-NA-PP1 0.5 µM addition does not reduce the intensity of Baz crescents compared to the previous cell cycle (loss of 5.3±24%, n=39). In*cdk1^as2^*neuroblasts, 1-NA-PP1 addition reduces the intensity of Baz crescents compared to the previous cell cycle (loss of 34.2±17%, n=36). 5 brains per condition across 3 experiments. **B**) Two consecutive divisions of cycling *baz^815-8^*, *cdk1^as2^* neuroblasts expressing Ba^Δ^z^LB^::GFP or Baz^ΔOD^::GFP, before and after exposure to 1-NA-PP1 0.5 µM. Scale bar: 5 µm. n=92 divisions in the presence of 1-NA-PP1 in 7 brains (Baz ^ΔLB^) and 75 divisions in 5 brains (Baz^ΔOD^), 2 experiments. **C**) Two consecutive divisions of cycling *cdk1^as2^* embryonic neuroblasts expressing Baz::GFP and His::RFP, before and after exposure to 1-NA-PP1 0.5 µM. **C’**) Ratio between the apical Baz signal at metaphase in the presence of 1-NA-PP1 0.5 µM and the Baz signal during the previous metaphase, in the absence of 1-NA-PP1 0.5 µM in *cdk1^as2^* embryonic neuroblasts. n=37 successive divisions in 31 neuroblasts across 5 embryos, 3 experiments. **D**) Cycling neuroblasts at metaphase. Scale bar: 5 µm. **D’**) Control cycling neuroblasts maintain stable Baz levels throughout metaphase (+0.4±6.1% in 75 seconds, n=52 neuroblasts, 4 brains) in the presence of 1-NA-PP1 0.5 µM. In CDK1 ^as2^ cycling neuroblasts, Baz levels decrease during metaphase in the presence 1-NA-PP1 0.5 µM (−9.8±7.3% in 75 seconds, n=49, 6 brains). 2 experiments. **E**) *cdk1^as2^* neuroblast first treated with 1-NA-PP1 0.5 µM for 1 hour, and then treated with Colcemid 50 µM. Neuroblasts stay polarized during metaphase (Baz crescents were maintained in 40/40 neuroblasts arrested in metaphase for at least 45’, 6 brains, 2 experiments). t0: NEB. Scale bar: 5 µm. **F**) *cdk1^as2^*neuroblast first treated with 1-NA-PP1 0.5 µM for 1 hour, then treated with Colcemid 50 µM for 30 minutes, after which 1-NA-PP1 was washed out (in this case 16.5’ after NEB). Some metaphase-arrested neuroblasts undergo Baz crescents coalescence following 1-NA-PP1 washout (n=19/51 neuroblasts, 7 brains, 3 experiments). t0: NEB. Scale bar: 5 µm**G**.) Cycling neuroblasts fixed at metaphase, exposed to 1-NA-PP1 0.5 µM, expressing Baz::GFP (green) and stained for aPKC (blue) and Miranda (magenta). Scale bar: 5 µm. **G’**) Basal Miranda and apical aPKC signals in metaphase neuroblasts exposed to 1-NA-PP1 0.5 µM. n=23 neuroblasts in 5 brains (*cdk1^+^*) and 27 neuroblasts in 4 brains (*cdk1^as2^*), 2 experiments. **H**) Cycling neuroblasts fixed at metaphase, exposed to 1-NA-PP1 0.5 µM, expressing Baz::GFP (green) and stained for Numb (magenta). Scale bar: 5 µ m**H**.**’**) Basal Numb signal in metaphase neuroblasts exposed to 1-NA-PP1 0.5 µM. n=55 neuroblasts in 6 brains (*cdk1^+^*) and 30 neuroblasts in 13 brains (*cdk1^as2^*), 2 experiments. **I**) In control neuroblasts, an apical Baz crescent (green) assembles in prophase and coalesces (green arrows) into a narrower, brighter crescent at NEB. Its intensity is maintained throughout metaphase. Upon partial inhibition of CDK1, an apical Baz crescent (green) assembles in prophase but fails to coalesce at NEB and its intensity decreases during metaphase. Basal polarity proteins (magenta) form more intense crescents than in controls.

### Partial inhibition of CDK1 destabilizes Baz crescents in cycling neuroblasts

Upon partial CDK1 inhibition, we also observed that the non-coalesced Baz crescents of cycling *cdk1^as2^* neuroblasts displayed a small but significant decrease in intensity comparing two consecutive time points 75 seconds apart during metaphase (−9.8±7.3%), whereas Baz crescent intensity in control neuroblasts was maintained (**Fig 2D, D’**), suggesting a role of CDK1 in maintaining Baz polarity. To investigate this possibility, we allowed neuroblasts to cycle into Colcemid-induced metaphase arrest under partial CDK1 inhibition. We reasoned that experimentally increasing the duration of metaphase would give enough time for unstable Baz crescents to completely disappear. However, Baz crescents remained stable in metaphase-arrested *cdk1^as2^*neuroblasts exposed to 0.5 µM 1-NA-PP1 (**Fig 2E**). The observation that non-coalesced crescents remain stable in metaphase enabled us to next test whether restoring normal CDK1 activity in metaphase-arrested neuroblasts could reinstate Baz crescent coalescence. Indeed, a 37% of metaphase-arrested neuroblasts with non-coalesced crescents gradually concentrated Baz in narrow and brighter crescents following 1-NA-PP1 washout (**Fig 2F**). Thus, partial inhibition of CDK1 can briefly destabilize Baz crescents in cycling neuroblasts, but not in metaphase-arrested neuroblasts. Furthermore, NEB and Baz coalescence can be uncoupled by temporally interfering with CDK1 activity.

### Partial inhibition of CDK1 leads to non-coalescence of apical aPKC crescents and to increased basal Numb levels

The function of Baz coalescence during neuroblast polarization is unknown. As one key function of Baz is the recruitment of aPKC, unsurprisingly, aPKC also localized in dimmer apical crescents upon partial CDK1 inhibition (**Fig 2G, G’**). As aPKC orchestrates basal cell fate determinant localization by directly phosphorylating them or their adapters, we tested whether weaker aPKC polarity was correlated with abnormal basal polarization. Indeed, Numb basal crescents were significantly (41±51%) more intense in *cdk1^as2^* neuroblasts compared to controls under partial CDK1 inhibition (**Fig 2H, H’**). Miranda crescents also displayed a slight, statistically insignificant increase (20±49%) in intensity upon CDK1 partial inhibition (**Fig 2G, G’**). In conclusion, partial inhibition of CDK1 prevents the coalescence of apical polarity crescents and leads to increased levels of basal fates determinants (**Fig 2I**).

### Baz is specifically phosphorylated in asymmetrically dividing cells of the central and peripheral nervous system on Serine180, a consensus CDK phosphorylation site

To determine whether the observed effects of CDK1 inhibition on Baz localization are direct, we interrogated the Baz amino acid sequence for full consensus CDK phosphorylation sites [ST]Px[KR] (Songyang et al., 1994). We identified three phosphorylation sites, among which two have been reported to be phosphorylated in phosphoproteomic approaches in *Drosophila*: Serine S180 (Bodenmiller et al., 2008, Zhai et al., 2008, Hilger et al., 2009) and Serine 417 (Hu et al., 2019). The presence of a Cyclin-binding motif ([RK]XL(X)[FYLIVMP], (Lowe et al., 2002) and of phospho-dependent Cks1-binding motifs ([FILPVWY]XTP, (McGrath et al., 2013) further suggests the possibility of CDK regulation of Baz (**Fig 3A**). Given its proximity to both a Cyclin-binding motif and a Cks1-binding motif, we focused our analysis on Serine 180 and generated a phosphospecific antibody (anti-Baz-pS180), which we first used on larval brains after trichloroacetic acid (TCA) fixation (Hayashi et al., 1999). Remarkably, a Baz-pS180 signal was only detected in dividing neuroblasts, and neither in interphase neuroblasts nor in neighboring neuroepithelial, regardless of their cell cycle stage (**Fig 3B, C**).

**Figure 3:**
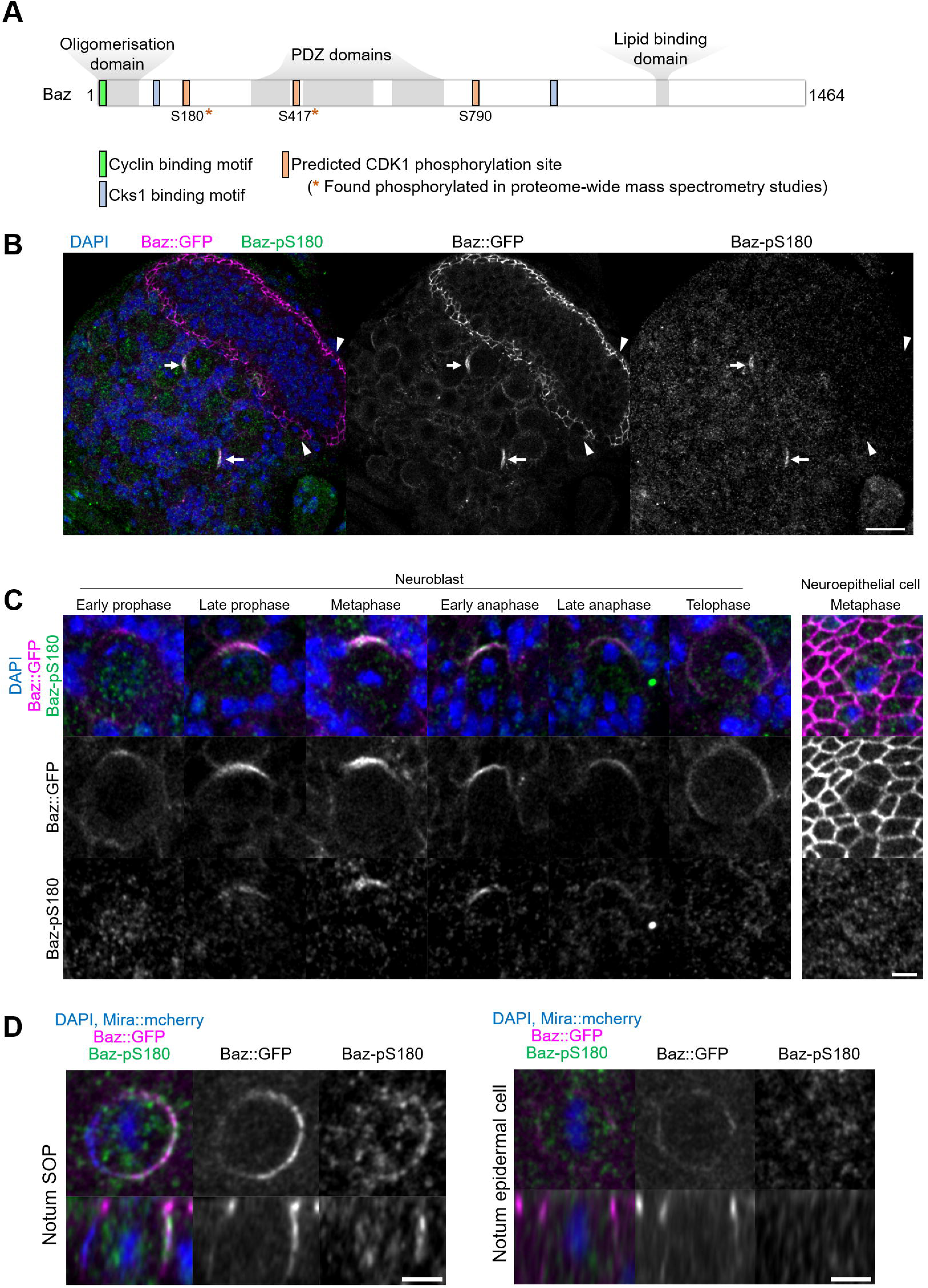
Asymmetric cell division-specific phosphorylation of Baz-S180. **A**) Structure of the Baz protein. Consensus motifs suggesting a potential phosphoregulation of Baz by CDK1 are highlighted. **B**) Fixed fly brain expressing Baz::GFP (magenta) and stained with a phospho-specific antibody against Baz-pS180 (green). Arrows: mitotic neuroblasts. Arrowheads: mitotic neuroepithelial cells. Scale bar: 20 µm. **C**) Left set of panels: cell cycle stages of fixed neuroblasts. Right set of panels: metaphase neuroepithelial cell. Scale bar: 5 µm. A Baz-pS180 signal was observed in 118/118 metaphase neuroblasts and 0/31 metaphase neuroepithelial cells (13 brains, 3 experiments). **D**) Left: notum microchaete sensory organ precursor in metaphase, at 16h APF. Right: notum epidermal cell in metaphase, at 16h APF. Bottom panels: orthogonal view. Scale bar: 5 µm.

This difference in Baz-S180 phosphorylation between asymmetrically dividing mesenchymal neuroblasts and symmetrically dividing neuroepithelial cells may arise from differences in cell division modes and/or cell type. To test this, we stained Baz-pS180 in the pupal notum at 16 hours after puparium formation (16h APF), where we could compare symmetric and asymmetric cell divisions of epithelial cells. Again, mitotic asymmetrically dividing sensory organ precursors (SOPs) were positive for Baz-pS180 whereas symmetrically dividing epidermal cells were not (**Fig 3D**). Similarly, in larval imaginal disks, dividing SOPs were positive for Baz-pS180 while symmetrically dividing columnar epithelial cells were not (**Fig S3A, B**). Interestingly, the CDK1 interacting partner CyclinA was recently found to localize to the apical-posterior cortex of dividing SOPs (Darnat et al., 2022). We confirmed that cortical CyclinA colocalizes with Baz-pS180 in mitotic SOPs (**Fig S3C**), potentially providing a mechanism for this asymmetric cell division-specific phosphorylation event. This prompted us to examine the localization of CDK1-associated Cyclins in neuroblasts. However, neither Cyclin A, Cyclin B nor Cyclin B3 localized to the cortex of dividing neuroblasts, indicating that neuroblast-specific phosphorylation of Baz-S180 is not achieved through cortical Cyclin localization in the larval brain (**Fig S3D**). We conclude that at least in the larval brain, larval imaginal discs and the pupal notum, Baz-S180 is specifically phosphorylated during asymmetric cell divisions.

### Phosphorylation of Baz-S180 affects the timing of neuroblast basal polarization

Having established that Baz-S180 is phosphorylated in vivo, we next tested its functional relevance in larval neuroblasts. To this end, we generated UASz-driven (DeLuca and Spradling, 2018) constructs containing either wildtype (Baz^WT^::GFP), non-phosphorylatable (Baz^S180A^::GFP) or phosphomimetic (Baz^S180D^::GFP) GFP-tagged versions of Baz. To assess the function of these constructs in the absence of endogenous Baz, we used UAS-driven RNAi against RFP to deplete a functional Baz::mScarlet-I CRISPR knock-in that we generated previously (Houssin et al., 2021). This resulted in an apparently complete depletion of Baz::mScarlet and expected Mira localization defects (Atwood et al., 2007, Atwood and Prehoda, 2009), which were rescued by expression of our Baz^WT^::GFP construct (**Fig 4A**). We proceeded to investigate the localization and function of our Baz-S180 phosphomutant constructs in Baz::mScarlet-I-depleted neuroblasts. First, we examined their localization in cycling neuroblasts and observed that both phosphomutant constructs seemingly localized similar to Baz^WT^::GFP throughout the cell cycle (**Fig 4B, C**). All constructs were likewise stably retained at the apical pole in metaphase-arrested neuroblasts (**Fig 4D**). We also took advantage of the Baz^S180A^::GFP construct to test the specificity of the Baz-pS180 antibody. In the absence of endogenous Baz, Baz^WT^::GFP apical crescents were positive for Baz-PS180 whereas Baz^S180A^::GFP apical crescents were not, confirming the specificity of the antibody (**Fig S3E**). We next assessed the functionality of Baz-S180 phosphomutants constructs in establishing basal polarity. In both phosphomutants, we detected a significant delay in the formation of basal Mira crescents. These crescents reached an equivalent cortical/cytoplasmic ratios to Baz^WT^::GFP-expressing neuroblasts only at metaphase in Baz^S180A^::GFP and anaphase in Baz^S180D^::GFP (**Fig 4E, F**). In conclusion, phosphorylation of Baz-S180 is not necessary for the coordination of Baz localization throughout the cell cycle but is involved in the timely establishment of basal polarity in larval neuroblasts.

**Figure 4:**
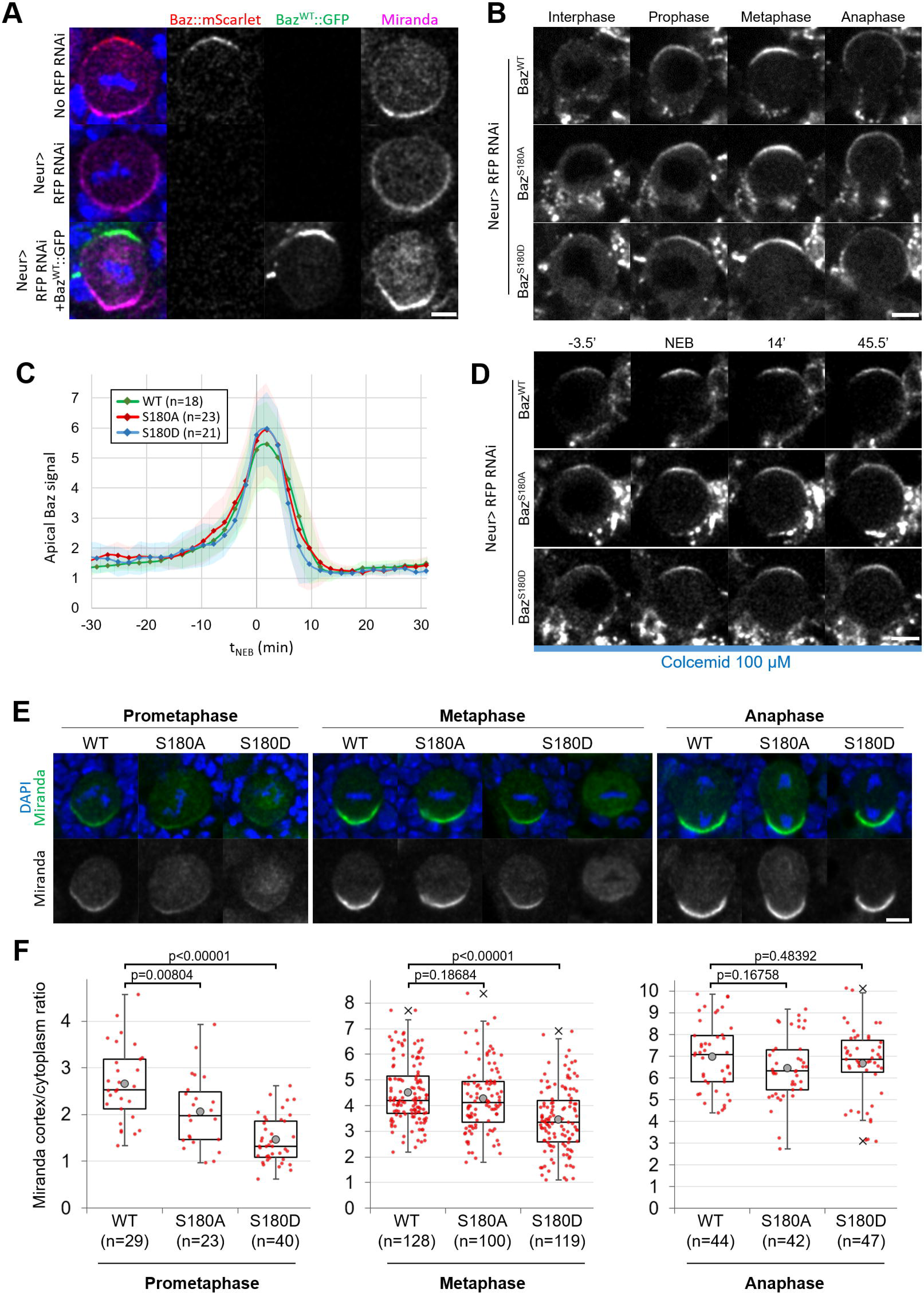
Localisation and function of Baz-S180 phosphomutants in neuroblasts. **A**) Fixed metaphase neuroblasts expressing Baz::mScarlet (red) and stained for Miranda (magenta), with or without Neur-GAL4 driving RFP RNAi and Baz^WT^::GFP (green) expression. Scale bar: 5 µm. **B**) Live neuroblasts depleted of endogenous Baz::mScarlet and expressing Baz^WT^::GFP, Baz^S180A^::GFP or Baz^S180D^::GFP. Scale bar: 5 µm. **C**) Intensity of the apical Baz::GFP signal in cycling neuroblasts depleted of endogenous Baz::mScarlet, normalized to the intensity of the cytoplasmic Baz signal in interphase. t0: NEB. n=18 divisions (Baz^WT^::GFP), 23 divisions (Baz^S180A^::GFP) and 21 divisions (Baz^S180D^::GFP). 5 brains for all conditions, 2 experiments. **D**) Live metaphase-arrested neuroblasts depleted of endogenous Baz::mScarlet and expressing Baz^WT^::GFP, Baz^S180A^::GFP or Baz^S180D^::GFP. T0: NEB. Scale bar: 5 µ m**E**.) Fixed neuroblasts depleted of endogenous Baz::mScarlet and expressing Baz^WT^::GFP, Baz^S180A^::GFP or Baz^S180D^::GFP, stained for Miranda (green) and DAPI (blue). Two cases are displayed for Baz^S180D^::GFP in metaphase, one showing polarized cortical Miranda (left) and the other mostly cytoplasmic Miranda (right). Scale bar: 5 µm. **F**) Ratio between the basal Miranda cortical signal and the cytoplasmic signal. n=29 prophases, 128 metaphases, 44 anaphases in 17 brains (Baz^WT^::GFP), 23 prophases, 100 metaphases, 42 anaphases in 18 brains (Baz^S180A^::GFP), and 40 prophases, 119 metaphases, 47 anaphases in 18 brains (Baz^S180D^::GFP). 2 experiments.

### Phosphorylation of Baz-S180 affects sensory organ formation

As Baz-S180 is also phosphorylated in mitotic SOPs (**Fig 3D**) we next analysed the functional relevance of this Baz phosphorylation in these cells. During asymmetric cell division of SOPs, cortical polarity controls the asymmetric segregation of Notch regulators, causing differential activation of the Notch pathway and thus the acquisition of different cell identities. The individual RNAi-mediated depletion of Baz or the Notch ligand Delta do not result in severe sensory organ formation defects. However, co-depletion of both results in a neurogenic phenotype and the near complete loss of sensory organs, indicative of a Notch loss of function (Houssin et al., 2021). We reproduced this observation by depleting both *delta* and endogenous *baz::mScarlet-I* (**Fig 5A**), which enabled functional analysis of GFP-tagged Baz-S180 phosphomutants in this context. Expression of our Baz::GFP constructs partially rescued the formation of macrochaetes, large sensory organs formed at the beginning of pupal development, but surprisingly not that of microchaetes, smaller sensory organs formed later during pupal development (**Fig 5B**). The Pannier-GAL4 driver is expressed in a wide stripe running across the entire length of the notum and scutellum, in which UAS-driven RNAi to knock down Baz::mScarlet appeared to be efficient, as mScarlet signal appeared to be entirely depleted in this zone. Unexpectedly, the expression pattern of our UASz-driven Baz::GFP constructs was largely restricted to two narrow posterior stripes, potentially explaining their inability to rescue the formation of more anterior microchaetes (**Fig 5C**). As microchaetes did not form in this context, we focused our analysis on macrochaetes.

**Figure 5:**
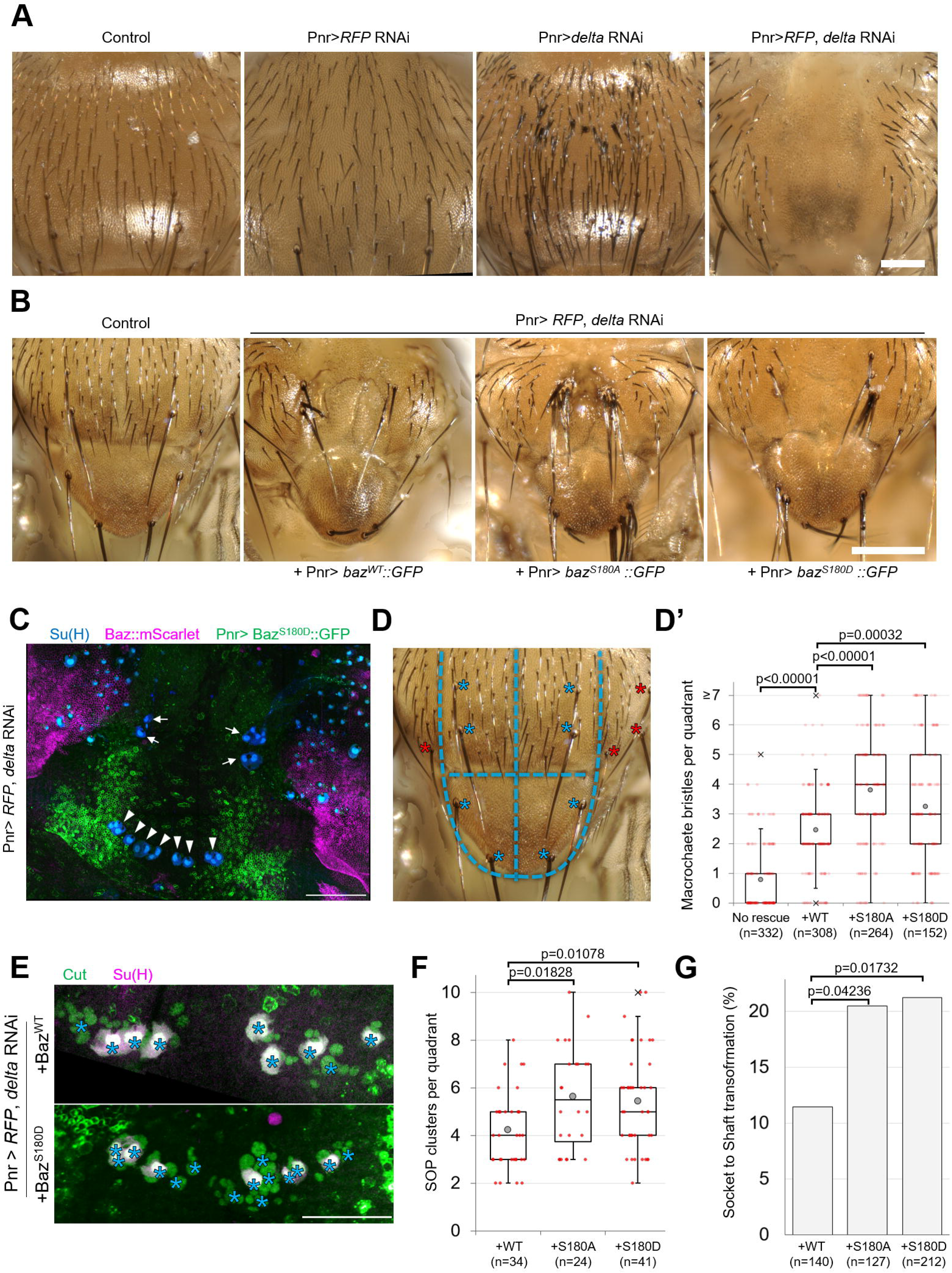
Baz-S180 phosphorylation regulates sensory organ formation. **A**) Adult nota with or without Pannier-GAL-driven RNAi of *baz::mScarlet* and/or *delta*. Scale bar: 100 µm. **B**) Control adult notum and *baz::mScarlet, delta* - depleted nota expressing Baz^WT^::GFP, Baz^S180A^::GFP or Baz^S180D^::GFP. Scale bar: 100 µm. **C**) Pupal notum stained for Su(H) (blue). Pannier-GAL4 drives both the depletion of endogenously expressed Baz::mScarlet and the expression of Baz^S180D^::GFP. Arrows: dorso-central macrochaetes. Arrowheads: scutellar macrochaetes. Scale bar: 100 µm. **D**) Quadrants defined for macrochaete bristles counting within the Pannier-GAL4 expression domain. The two upper quadrants include dorso-central macrochaetes and the two lower quadrants include scutellar macrochaetes (blues asterisks). Other macrochaetes (red asterisks) outside of the Pannier-GAL4 expression domain the were ignored. **D’**) Number of individual bristles per quadrant in *baz::mScarlet, delta* - depleted adult nota, with or without expression of Baz::GFP transgenes. n=332 quadrants in 83 flies (no rescue), 308 quadrants in 77 flies (Baz^WT^::GFP), 264 quadrants in 66 flies (Baz^S180A^::GFP) and 152 quadrants in 38 flies (Baz^S180D^::GFP). **E**) Pupal notum stained for Cut (green) and Su(H) (magenta). Asterisks: individual SOP clusters. Scale bar: 50 µm. **F**) Number of individual SOP clusters per quadrant in *baz::mScarlet, delta* - depleted pupal nota expressing Baz::GFP transgenes. N=34 quadrants in 9 nota (Baz^WT^::GFP), 24 quadrants in 6 nota (Baz^S180A^::GFP), and 41 quadrants in 11 nota (Baz^S180D^::GFP). 4 experiments. **G**) Percentage of cases of Socket cell to Shaft cell transformation cases in *baz::mScarlet, delta*-depleted pupal nota expressing Baz::GFP transgenes. n=140 SOP clusters in 9 nota (Baz^WT^::GFP), 127 SOP clusters in 6 nota (Baz^S180A^::GFP), and 212 SOP clusters in 11 nota (Baz^S180D^::GFP). 4 experiments. Statistical test: two-tailed Z score calculation of population proportions.

We observed that significantly more macrochaete bristles formed when expressing either phosphomutant compared to the rescue with Baz^WT^::GFP (**Fig 5D**). This excess of bristles could be indicative of defects in lateral inhibition, a Notch-dependent mechanism restricting the number of sensory organ precursors. It could also result from defects in asymmetric cell divisions in the sensory organ precursor lineage leading to cell fate transformations such as socket-to-shaft transformations. To investigate both possibilities, we stained pupal nota for SOP lineage (Cut) and socket cell (Su(H)) markers (**Fig 5E**). Rescue by the Baz-S180 phosphomutants in the *baz, delta* double RNAi background resulted in significantly more SOPs to be formed (**Fig 5F**) and in significantly more socket-to-shaft transformations (**Fig 5G**) than rescue by Baz^WT^::GFP, suggesting with a Notch loss of function during lateral inhibition and following asymmetric cell divisions, respectively. We conclude from this result that, in the absence of Delta, phosphorylation of Baz-S180 affects sensory organ formation both during the initial specification of SOPs, and during the asymmetric cell divisions they undergo.

### Human PARD3 is a substrate of CDK1/CyclinB1 in vitro

Given that we found the consensus CDK phosphosite S180 in Baz to be phosphorylated and functionally relevant in neuroblasts and SOP lineage formation, we wondered whether it is conserved in humans. Interestingly, although the sequence of the disordered region surrounding S180 does not appear to be conserved in humans, we identified on PARD3 a full consensus CDK phosphosite (S187) in close proximity, like S180 in*Drosophila* Baz, to a 14-3-3 binding site (**Fig 6A**). Previously, PARD3-S187 was found phosphorylated in high-throughput mass spectrometry studies (Lin et al., 2021). To test whether PARD3-S187 or other residues could be phosphorylated by CDK1 in humans, we purified full length human PARD3 from insect cells and used targeted mass spectrometry to identify PARD3 Ser/Thr phosphorylation in the presence or absence of CDK1/CyclinB1 complexes. Incubating PARD3 with CDK1 increased the detection of phospho Serines and Threonines. Unfortunately, we could not determine whether PARD3-S187 is phosphorylated *in vitro* by CDK1 because no peptides covering the region surrounding PARD3-S187 were confidently identified following Trypsin digestion. Nonetheless, other phosphopeptides containing consensus CDK phosphosites were detected. S728, a full consensus CDK phosphosite ([ST]Px[KR], (Songyang et al., 1994) was phosphorylated in the presence and absence of CDK1/CyclinB1. S1136 and S1332, minimal consensus CDK phosphosites ([ST]P], (Suzuki et al., 2015) were phosphorylated only in the presence of CDK1/CyclinB1 (**Fig 6B-D, Supplementary table 1**). In conclusion, these data argue that human PARD3 may be a direct substrate for CDK1/CyclinB1.

**Figure 6:**
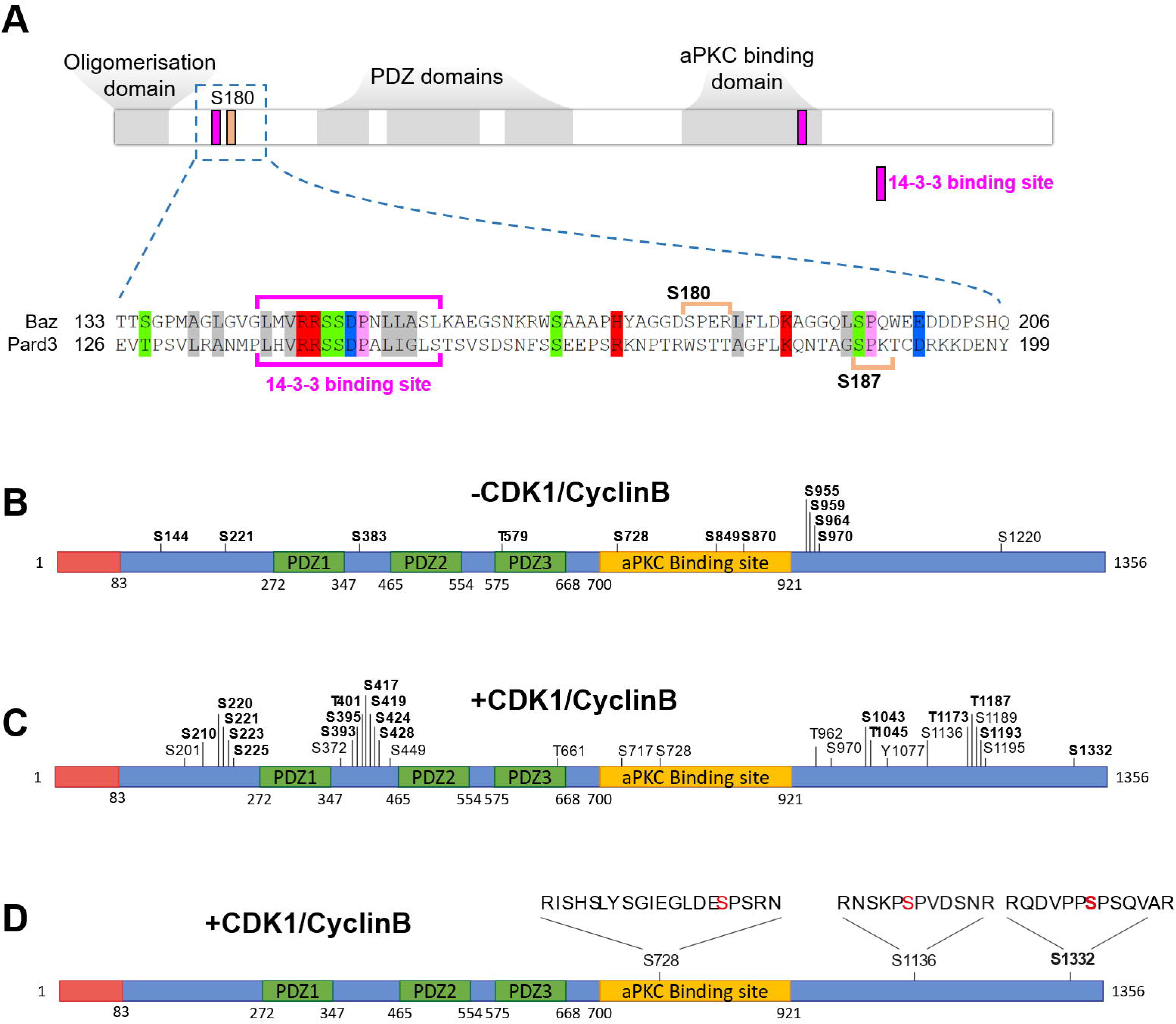
Human PARD3 is a substrate of CDK1 *in vitro*. **A**) Domains of PAR-3 proteins (top) and alignment of the regions surrounding the N-terminal 14-3-3 binding sites of *Drosophila* Baz and human PARD3. Magenta brackets: 14-3-3 binding site. Orange brackets: consensus CDK phosphorylation site sequence. **B-D**) Schematic representation of recombinant human PARD3 with Serine/Threonine phosphosites detected by mass spectrometry in the absence (**B**) or presence (**C,D**) of CDK1/CyclinB1. Sites in bold were detected in both replicates. In **D**, consensus CDK phosphorylation sites phosphorylated in the presence of CDK1/CyclinB1 are displayed with their peptide sequence.

## Discussion

### Partial inhibition of CDK1 does not abolish neuroblast polarity

Here, we first reinvestigated the role of CDK1 during neuroblasts asymmetric cell division. Based on the observation on fixed tissues that dominant negative or thermosensitive *cdk1* alleles do not prevent embryonic neuroblasts from cycling but disrupt their polarity, asymmetric cell division was proposed to strictly rely on high levels of CDK1 activity (Tio et al., 2001). However, using live imaging analysis and chemical genetics, we did not reproduce these observations: the apical polarity protein Baz still polarizes upon partial inhibition of CDK1 using an analog-sensitive allele of *cdk1*, both in embryonic and larval neuroblasts (**Fig 2**). A possible cause for this discrepancy could be that loss of Baz polarity was previously only reported with the dominant negative *cdk1^E51Q^* allele, for which neomorphic activity cannot be ruled out, whereas loss of polarity in the thermosensitive *cdk1* genetic situation was only reported for another apical polarity protein, Inscuteable. Considering that Baz functions upstream of Inscuteable (Schober et al., 1999), it is possible that Baz localization is not affected in this situation.

### High CDK1 activity triggers apical coalescence

While partial inhibition using the analog-sensitive allele of cdk1 did not abolish Baz polarity, it did prevent apical coalescence: large apical crescents formed during prophase failed to coalesce into smaller crescents at NEB (**Fig 2**). Although our observations show that high CDK1 activity is required for apical coalescence, the exact mechanism regulating this process remains elusive. Our previous observation that aPKC inhibition can result in apical crescents coalescing in the wrong direction (Hannaford et al., 2019) suggests that apical polarity proteins are involved in some aspects of this process. As coalescence was not affected in Baz-S180 phosphomutants (**Fig 4**), CDK1-dependent control of apical coalescence might be controlled by other CDK1 phosphosites on Baz or other CDK1 targets. RhoGAPs and RhoGEFs are promising candidates for this: first, apical coalescence of neuroblasts is driven by basal-to-apical actomyosin-driven cortical flows (Oon and Prehoda, 2019), reminiscent of the ones polarising *C. elegans* embryos (Munro et al., 2004) induced by asymmetric localization of the RhoGEF ECT-2 (Motegi and Sugimoto, 2006); second, ECT-2 can be phosphorylated by CDK1 (Niiya et al., 2005). Based on the observation that some RhoGEFs such as the fly ECT-2 homolog Pebble can be sequestered in the nucleus until NEB and that apical coalescence occurs at NEB, a tempting model for the temporal regulation of coalescence would be that the release of RhoGEFs at NEB triggers a basal-to-apical flow. However, we showed that NEB and apical coalescence can be temporally uncoupled by first partially inhibiting CDK1, then restoring its full activity after NEB has occurred (**Fig 2F**). Thus, although an involvement of NEB cannot be ruled out in its temporal control, it is ultimately high CDK1 activity that triggers apical coalescence. Interestingly, enlarged Baz crescents were observed in *aurora A* mutants (Wang et al., 2006). Given that CDK1 activates Aurora A (Van Horn et al., 2010), this raises the possibility that CDK1 drives apical coalescence *via* Aurora A activation.

### Function of apical coalescence

Beyond its regulation, the function of apical coalescence is unknown. Based on our observation that partial inhibition of CDK1 not only leads to defective coalescence but also to crescent instability in metaphase in cycling neuroblasts (**Fig 2D**), we initially speculated that coalescence, by concentrating apical polarity proteins, might reinforce positive feedback loops for apical protein concentration and thus stabilize them. However, partial inhibition of CDK1 and the resulting failure to coalesce surprisingly did not seem to affect crescent stability in metaphase-arrested neuroblasts (**Fig 2E**). A possible explanation for this difference between cycling and arrested cells could be that, in cycling cells, partial inhibition of CDK1 somehow prematurely triggers some anaphase events involving Baz crescents disassembly, uncoupling them from other anaphase events such as DNA segregation and cortical furrowing. Regardless of the reason for this difference, apical coalescence is not necessary for apical polarity maintenance. However, we did observe that basal Numb crescents significantly increased in intensity upon partial inhibition of CDK1 (**Fig 2C, C’**). We speculate that enlarged apical crescents might be able to exclude basal protein further down the apico-basal axis, concentrating basal components in smaller basal crescents. Brighter Numb crescents did not appear noticeably smaller than in controls in our fixed tissue analysis, but live imaging of neuroblasts in culture would be necessary to accurately measure basal crescents size. Nonetheless, aPKC activity is sufficient to exclude basal fate determinants from the apical and lateral regions even when coalescence fails. An alternative explanation could be that, upon partial inhibition of CDK1, a longer cell cycle allows neuroblast to synthesize more Numb, raising the interesting possibility of a modulation of the strength of basal asymmetry by cell cycle duration. Finding ways of interfering with coalescence without increasing cell cycle length will be necessary to further study the functional role. Finally, failed apical coalescence indicative of loss of basal-to-apical flows could affect the properties of the basal cortex, impacting basal fate determinants localization.

### Asymmetric cell division-specific phosphorylation of Baz

Remarkably, we show that Serine 180 of Baz is phosphorylated specifically in mitotic cells undergoing asymmetric cell division, and not in their symmetrically dividing neighboring cells in the central and peripheral nervous system (**Fig 3**). To the best of our knowledge, this is the first description of an asymmetric cell division-specific phosphorylation event in the *Drosophila* central and peripheral nervous system. How is it achieved? Our interest in Baz-S180 was originally prompted by the fact that it is a consensus CDK phosphorylation site, and by its proximity to Cyclin and Cks1-binding sites (**Fig 3A**). However, CDK1 is also involved in symmetric cell division. Asymmetric cell division-specific phosphorylation of Baz by CDK1 might result from higher CDK1 activity than in symmetrically dividing cells, from an asymmetric cell division-specific adaptor between CDK1 and Baz or from asymmetric cell division-specific inhibition of Baz-S180 dephosphorylation. It is also noteworthy that Cyclin A colocalizes with Baz in asymmetrically dividing SOPs (Darnat et al., 2022) (**Fig S3D**), perhaps explaining SOP-specific Baz-S180 phosphorylation. However, none of the CDK1-associated Cyclins localize asymmetrically to the mitotic neuroblasts cortex (**Fig S3C**). Alternately, Baz-S180 could simply be phosphorylated by another as-yet-unknown kinase specifically expressed in asymmetrically dividing cell and only active during mitosis.

### Phosphorylation of human PARD3 by CDK1

Wondering whether Baz-S180 was conserved in humans, we identified in silico four consensus CDK phosphorylation sites on human PARD3. One of them, PARD3-S187, is like Baz-S180 in close proximity to an N-terminal 14-3-3 binding site (**Fig 6A**) regulating PAR-3 proteins localization (Benton and St Johnston, 2003). We could not determine whether PARD3-S187 is phosphorylated by CDK1 in our *in vitro* experiment with purified proteins (**Fig 6B-D**), but a novel *in vitro* phosphorylation assay on fixed lymphoblasts did not identify PARD3-S187 as a CDK1 target (Al-Rawi et al., 2023), although CDK1 phosphorylated other PARD3 sites in both assays. Thus, either CDK1 does not regulate PARD3-S187 (which, as discussed above, is also possible for Baz-S180), or *in vitro* assays do not recapitulate what makes asymmetric cell division-specific phosphorylation of Baz-S180 possible. Thus, the potential phosphoregulation and role of PARD3-S187 and other CDK consensus phosphorylation sites may need to be investigated in *in vivo* contexts such as the PARD3-regulated asymmetric cell division of radial glia in mice (Bultje et al., 2009). Interestingly, Cks1 stimulates non-proline directed phosphorylation of CDK1 targets in humans (Al-Rawi et al., 2023). If this property of Cks1 is conserved in *Drosophila*, the presence of a Cks1 binding site on Baz (**Fig 3A**) opens the possibility that the potential regulation of Baz by CDK1 may not be limited to its consensus Proline-directed phosphosites.

### Function of Baz-S180 phosphorylation in neuroblasts

We used Baz-S180 phosphomutants in the absence of endogenous Baz to investigate the function of this phosphorylation, expecting it to be necessary for the timing of Baz polarization based on its mitosis-specificity, and/or for Baz ability to polarize, given its asymmetric cell division-specificity. However, neither polarization itself, nor its timing, nor its maintenance were affected in Baz-S180 phosphomutant neuroblasts (**Fig 4B-D**). Thus, Baz-S180 phosphorylation is not the temporal cue coordinating cell polarity and the cell cycle, and the observed effects upon full or partial CDK1 inhibition on Baz localization and dynamics are likely caused by other phosphosites on Baz and/or other CDK1 substrates. Although apical Baz crescents were apparently unaffected in Baz-S180 phosphomutants, basal Miranda crescents appeared weaker, sometimes barely detectable, in prometaphase and metaphase. In contrast, they were indistinguishable from controls in anaphase (**Fig 4E-F**), perhaps through the “telophase rescue” mechanism that somehow restores defective basal polarity in *baz* and *insc* mutant neuroblasts (Schober et al., 1999, Wodarz et al., 1999). Why the establishment of basal polarity is delayed in Baz-S180 phosphomutants remains unclear. As the function of the PAR complex is to exclude Miranda from the apical pole by phosphorylating it, defective recruitment of Miranda to the cortex could be explained by a gain of function of the PAR complex in Baz-S180 phosphomutants. We speculate that it might be caused by an excessive transfer of aPKC from an immobile apical Baz/aPKC/PAR-6 complex to a more diffusible Cdc42/aPKC/PAR-6 complex (Rodriguez et al., 2017), able to phosphorylate Miranda further down the apico-basal axis.

### Function of Baz-S180 phosphorylation in sensory organs formation

We also showed that sensory organs formation is affected in Baz-S180 phosphomutants: we observed an excess of bristles (**Fig 5D’**) caused by the specification of too many sensory organ precursor cells (**Fig 5F**) and because of lineage transformations leading to an excess of shaft cells (**Fig 5G**). The latter is not surprising: shaft-to-socket transformations are a well-studied phenotype caused by a failure to activate the Notch pathway following one of the asymmetric cell divisions taking place in the SOP lineage (Schweisguth, 2015), in which we observed asymmetric cell division-specific phosphorylation of Baz-S180, and are likely due to defective Baz-regulated asymmetric segregation of Notch regulators and/or Baz-dependent assembly of Notch clusters (Houssin et al., 2021). In contrast, the specification of an excess of SOPs is at first glance more puzzling. Why would the lateral inhibition mechanism specifying SOPs be regulated by a phosphorylation event that appears to be restricted to asymmetric cell division? It is possible that S180 has a phosphorylation-independent function in lateral inhibition, or that S180 phosphorylation is actually not restricted to asymmetric cell division when lateral inhibition takes place. However, the concept of Baz-S180 phosphorylation being indeed asymmetric cell division-specific and the possibility that Baz-S180 phosphorylation regulates lateral inhibition are not mutually exclusive given the following observation: macrochaete SOPs arise from clusters of G2-arrested cells (Usui and Kimura, 1992). Indeed, CDK1 (or other cell cycle regulated kinases) activity in G2 (Hochegger et al., 2008) could be sufficient to phosphorylate Baz-S180, and this phosphorylation event could arguably be considered to be asymmetric cell division-specific, as it would occur during the G2 phase of cells that all have the potential to perform asymmetric cell division (until this potential is later restricted to one SOP by lateral inhibition). Regardless of these considerations, as lateral inhibition is Notch-dependent (Simpson, 1990), it could be regulated by Baz through recently described mechanisms outside of its “canonical” role in asymmetric cell division: Baz recruits Notch at the interface of daughter cells following asymmetric cell division (Wu et al., 2023b, Houssin et al., 2021) and is somehow involved in the activation of the Notch intracellular domain following its cleavage (Wu et al., 2023a). Finally, it is noteworthy that Baz-S180 non-phosphorylatable and phosphomimetic mutants give the same phenotypes (**Fig 4**, **5**) (with the exception of basal polarization of Miranda being more delayed in *baz^S180D^* than in *baz^S180A^*in neuroblasts), likely because phosphomimetics do not always faithfully reproduce the effects of a phosphorylation (Gogl et al., 2021).

### Robustness of Baz localization and function

Across the few decades during which Baz has been studied, a recurring observation is that Baz-dependent processes are often robust, involving context-specific functional redundancies between Baz and other proteins. Therefore, despite the central importance of Baz throughout development, disrupting Baz function alone can sometimes lead to mild phenotypes which are significantly worsened by concomitantly interfering with other proteins. Accordingly, Baz functions redundantly with the polarity protein PAR-6 in tracheal terminal cells branching (Jones and Metzstein, 2011), with the transmembrane protein Crumbs in the maintenance of epithelial polarity (Müller and Wieschaus, 1996), with the Notch ligand Delta in the formation of sensory organs (Houssin et al., 2021), and with the apical polarity Pins in the generation of size asymmetry during neuroblasts asymmetric cell division (Cai et al., 2003). Beyond redundancies with other proteins, the robustness of Baz function relies on a lipid binding domain and an oligomerization domain, which are both sufficient on their own to support Baz localization and function (Kullmann and Krahn, 2018). In this light, it is perhaps not surprising that the phenotypes that we describe in neuroblasts remain relatively mild: partial inhibition of CDK1 does not abolish apical nor basal polarity (**Fig 2**), and Baz-S180 phosphomutants display no localization defect and only a delay in basal polarity establishment (**Fig 3**). It is conceivable that, in these situations, Baz localization and/or function is rescued by another protein. A very likely candidate for this is Pins, known to function redundantly with Baz in neuroblasts (Cai et al., 2003, Yu et al., 2000, Izumi et al., 2004). Could Pins be phosphorylated by mitotic kinases, and in this case, would both Baz and Pins need to lose the temporal cue provided by phosphorylation to uncouple cell polarity from the cell cycle? In the absence of Pins, would apical coalescence become necessary for the maintenance of apical polarity? Would Baz-S180 phosphomutants fail to polarize? Addressing these questions will be the focus of further studies.

## Supplementary figure legends

**Supplementary Figure 2 – related to Figure 2**. Cumulative sum chart of NEB events in *cdk1^as2^*brains, normalized to the total number by the time of 1-NA-PP1 addition (t0). N=141 divisions in 4 brains (0.5 µM), 77 divisions in 4 brains (1 µM), and 100 divisions in 3 brains (2 µM), 1 experiment.

**Supplementary Figure 3 – related to Figure 3. A**) Leg disk sensory organ precursor in metaphase, at 0h APF. Bottom panels: orthogonal view. Scale bar: 5 µm. **B**) Wing disk columnar epithelium cell in metaphase. Bottom panels: orthogonal view. Scale bar: 5 µm. **C**) metaphase SOP expressing Pon::RFP (red) under the control of Neur-GAL4, stained for CycA (magenta) and Baz-pS180 (green). **D**) Left panels: fixed larval brain neuroblasts expressing Baz::GFP (magenta) and stained for Cyclin A (green). Middle and right panels: live larval brain neuroblasts expressing endogenously expressed CycB3::GFP or CycB::GFP protein traps. E) Fixed neuroblasts depleted of endogenous Baz::mScarlet and expressing Baz^WT^::GFP or Baz^S180A^::GFP. Scale bar: 5 µm.

**Supplementary Table 1. Confidently identified phosphopeptides from phospho-proteomics of human PARD3 in the presence or absence of CDK1/CyclinB1. Sheet 1**: All confidently identified phosphopeptides, listed by sample condition and replicate. N=2; “no kinase” and “kinase” reactions. Obs. Mass, observed mass; Calc. mass, calculated mass. Red and lower-case s, t, are the site ID from Proteome Discoverer PhosphoRS3.1 where the phosphoRS Site Probability is >90%. **Sheet 2** : All confidently identified phosphopeptides annotated and arranged according to phosphosite location and whether identified in one or both conditions (no kinase/kinase) and replicates. **Sheet 3** : Summary list of all confidently identified phosphopeptides annotated and arranged according to phosphosite location and surrounding peptide motif to identify conserved CDK1 proline-directed phosphosites. **Sheet 4**: Summary list filtered for phosphopeptides identified only in “kinase” reactions, arranged by motif surrounding the identified phosphosites (as in Sheet 3).

**Supplementary Table 2: genotypes.**

**Supplementary Table 3: origin of stocks.**

## Materials and methods

### Fly stocks and genetics

Flies were reared on standard corn meal food at 251l°C, except for RNAi-expressing larvae and their corresponding controls (**Fig 4, 5, S3E**), which were placed at 291l°C from the L1 larval stage to the L3 larval or pupal stage, at which point they were dissected, or the adult stage, at which point they were observed under anesthesia with CO _2_. For the genotypes of the animals used in each experiment, see **Supplementary Table 1**. For the origins of the stocks used, see **Supplementary Table 2**.

### Immunostainings

The Baz-PS180 antibody was raised in sheep that were injected with YAGGDS*PERLF – (where S* is phospho Serine). The fourth bleed was affinity purified using negative selection against a YAGGDSPERLF peptide and positive selection using the YAGGDS*PERLF phospho-peptide. The polyclonal Numb antibody was generated in sheep using GST-full length Numb protein (CG3779-PA) and affinity purified. Specificity was tested by staining mitotic *numb^15^*(Le Borgne and Schweisguth, 2003) mutant SOP clones in the notum, where immune reactivity dropped to background levels. Polyclonal anti Mira antibody was raised in rabbit using the CSPPQKQVLKARNI peptide and affinity purified. Specificity was tested in mitotic *mira^KO^ (Ramat et al., 2017)* larval neuroblast clones where immune reactivity dropped to background levels. Polyclonal anti Mira antibody raised in sheep was generated using a RLFRTPSLPQRLR peptide followed by affinity purification.

For all immunostainings except for Baz-PS180 phosphostainings, larval brains, larval imaginal disks or pupal nota were dissected in phosphate buffered saline (PBS) and fixed in 4% Formaldehyde (Sigma F8775) for 20 minutes at room temperature (RT). For Baz-PS180 phosphostainings (**Fig 3, S3**), tissues dissected in PBS and fixed in 2% Trichloroacetic acid for 3 minutes at RT. Regardless of the fixation procedure, tissues were then permeabilised in for 1 hour in PBS-Triton 0.1% (PBT) at RT, and incubated overnight at 4°C in a solution of primary antibody diluted in PBT. They were rinsed twice and washed for 1 hour in PBT, incubated 1 hour at RT in a solution of DAPI 1/1000 and fluorophore-coupled secondary antibodies (Thermo Fisher) diluted in PBT, rinsed twice and washed in PBT, rinsed twice in PBS, rinsed in 50% glycerol (Sigma 49781) and finally mounted between glass and coverslip in Vectashield (Vector Laboratories H-1000). Primary antibodies used were as follows: rabbit anti-PKCζ (Santa Cruz, sc-17781, 1:500); Sheep-anti-Mira (1:1000); Rabbit anti Mira; Sheep anti-Numb (1:500); Mouse anti-Cyclin A (DSHB A12 1/500); Mouse anti-Su(H) (Santa Cruz sc-398453 AF647, 1/1000); Mouse anti-Cut (DSHB 2B10 1/200), Sheep anti-Baz-pS180 (this study, 1/200).

### Live imaging of larval brains

Every reference to Schneider’s medium corresponds to glucose-supplemented (1g1ll^−1^) Schneider’s medium (SLS-04–351Q). Tissue mounting and live imaging were performed as described (Januschke and Loyer, 2020). Entire brains were dissected from L3 larvae in Schneider’s medium and isolated from the surrounding imaginal discs. Particular care was taken to avoid pulling on brains at any time during the dissection and damaged brains were discarded. Isolated brains were transferred to a drop of Fibrinogen dissolved in Schneider’s medium (101lmg/ml) on a 231lmm Glass bottom dish (WPI), which was then clotted by addition of Thrombin (1001lU/ml, Sigma T7513). Clots were then covered in Schneider’s medium, taking into account the existing volumes of Schneider+Fibrinogen and Thrombin forming the clots for a final volume of 200 µl. For experiments involving 1-NA-PP1 addition, control and analog-sensitive tissues were mounted on different clots in the same dish and 1-NA-PP1 was added during imaging by adding 200 µl of Schneider with double the desired final concentration of 1-NA-PP1 (*e.g.* 200 µl of 20 µM 1-NAPP-1 in Schneider medium were added to the coverslip for a final concentration of 10 µM). Where applicable, neuroblasts were arrested in metaphase by exposure to 100 µM Colcemid (Sigma 234109).

### Live imaging of embryonic neuroblasts

Embryos were collected 3 to 4 hours after egg laying, manually dechorionated by rolling them on double-sided tape and transferred to a well containing Schneider’s medium. The posterior tip of the eggshell was nicked with a sharp forceps and the embryos were squeezed out by pressing from the anterior to the posterior of the eggshell. The mounting in Fibrin clots, live imaging and exposure to 1-NA-PP1 of de-shelled embryos then proceeded as described above for larval brains, except for the imaging medium, this time supplemented with glucose (11lg1ll−1), Insulin (75 µg/ml, Lonza, BE02-033E20), 10% Fetal Calf Serum and 2.5% fly extract (Drosophila Genomics Resource Center, 1645670).

### Cloning and transgenesis

The analog sensitive allele *cdk1^as2^* was generated by genome editing using the guide RNA GATCTTTGAATTCCTATCGA TGG and a template introducing the F80A (TTT to GCC) mutation and a silent mutation at L83 (CTA to TTG) to prevent re-editing by the guide RNA. The UASz-driven GFP-tagged Baz transgenes correspond to the Baz-PA isoform, in which 6xHiseGFP was introduced at the same position (between K40 and P41) and with the same linker peptides as the functional Baz::GFP protein trap (Buszczak et al., 2007) and Baz::mScarlet-I knock-in (Houssin, Pinot et al. 2021) using Gibson assembly cloning into UASz 1.0 (DGRC 1431). To generate the non-phosphorylatable and phosphomimetic constructs in the relevant Serines in Baz, site-directed mutagenesis was used to introduced alanine (GCG) or aspartic acid (GAG), respectively. All vectors were verified by sequencing (vector sequences available upon request). Transgenic lines were generated by site-directed transgenesis (https://www.flyfacility.gen.cam.ac.uk/Services/Microinjectionservice) into attP2.

### Image processing and analysis

All images were processed and filtered using ImageJ. For all images, we applied a 3D gaussian blur with X, Y and Z sigmas of 0.8 pixels. For all images where applicable, we applied a Gamma filter with a value of 0.5 on the DAPI or Histone::RFP channels. For the timing of NEB, we considered that NEB has started as soon as the Baz signal is no longer excluded from the nucleus. Measurement of cortical signal intensity was performed as described before using our custom rotating linescans macro (Januschke and Loyer, 2020). Cytoplasmic signal intensity was measured inside polygonal shapes excluding the cortex, the nucleus or chromosomes. The background fluorescence was measured outside of brains and subtracted from any measured signal. P values displayed over boxplots were calculated using a non-parametric two-tailed Mann–Whitney U test, except for Fig 3G, which corresponds to a two-tailed Z score calculation of population proportions.

### Purification of human PARD3, kinase assay and mass spectrometry

4 litre SF9 cell pellet from 6xHis-MBP-hsPARD3 was thawed on ice and topped up to 80 mL with Par3 Buffer (20mM HEPES, 250mM NaCl, 0.5mM TCEP, 10% Glycerol, pH 7.4) and cOmplete, EDTA-free Protease inhibitor cocktail (Roche) and 10 µl Benzonase (Merck, 70746) were added. Samples were homogenised 5x on ice in a 40 mL Dounce Homogeniser. Samples were centrifuged at 60,000 x g at 4°C for 1 hour. 5 mL amylose resin (MRC Reagents and Services) was pre-equilibrated in Par3 Buffer. Supernatant was syringe filtered through a 0.45 μm and supernatant was added to 5 mL of pre-equilibrated resin. Samples were then incubated on a rotary spinner at 4°C for 2 hours. Samples were centrifuged at 500 x g at 4°C for 5 minutes, and the flow-through was removed. 40ml of buffer was added to the resin. The first wash step included 0.1% CHAPS and protease inhibitor as previously. During this wash, sample was incubated on a rotary spinner at 4°C for 30 minutes. It was then centrifuged at 500 xg, 4°C for 1 minute and the excess buffer was poured off. The resin was subsequently washed briefly in Par3 buffer twice. The full-length Par3 protein was cleaved from MBP while on resin overnight by addition of 100 uL of HRV-3C protease (homemade) in 10 mL of Par3 Buffer. Samples were analysed by SDS-PAGE and Coomassie gel stain.

Elutions from the affinity chromatography step were diluted 1:5 in Buffer A (20mM HEPES, 0.5mM TCEP, 10% Glycerol) and loaded onto a 1 mL HiTrap Q column (Fisher) pre-equilibrated in Buffer A. Proteins were eluted from the column over the course of an increasing NaCl gradient from 0-100% Buffer B (20mM HEPES, 1M NaCl, 0.5mM TCEP, 10% Glycerol, pH7.4) across 10 column volumes, and peak fractions were collected. Samples from multiple peak fractions were analysed by SDS-PAGE and Coomassie gel stain. The fractions containing purified protein were collected.

Combined samples concentrated to 1 mL from anion exchange were injected onto a Superose 6 Increase 10/300 GL sizing column (Cytiva) pre-equilibrated in Par3 Buffer and peak fractions were collected and analysed by SDS-PAGE and Coomassie gel stain. Protein fractions were identified, aliquoted snap frozen using LQN_2_ and stored at −80°C.

### In vitro kinase phosphorylation assay

1 µg purified recombinant human PARD3 was incubated with 750ng of GST-CDK1/CyclinB1 (abcam; ab271456) in kinase buffer (50 mM Tris-HCl [pH 7.5], 0.1mM EGTA, 10mM MgCl2, 2mM Dithiothreitol [DTT]) in the presence of 2mM ATP. Reactions were incubated for 30 minutes at 30°C, shaking at 1000 rpm, and stopped by boiling for 5 mins with LDS/RA loading buffer.

### In-gel digestion

For phospho-site identification, kinase reactions were separated via SDS-PAGE electrophoresis, stained with Coomassie blue and gel pieces subjected to an in-gel digestion. Bands with corresponding protein were cut and washed sequentially in water, 50% acetonitrile (ACN), 0.1 M NH4HCO3, and 50% ACN/ 50 mM NH4HCO3 (all from Sigma-Aldrich) for 10 mins at RT, shaking at 1000 rpm. Proteins were alkylated with 10 mM DTT/0.1 M NH4HCO3 at 37°C for 20 mins and 50 mM iodoacetamide/ 0.1 M NH4HCO3 for 20 mins. Samples were washed as above. Gel pieces were shrunk with ACN for 15 mins, supernatant removed, and gel pieces dried using Speed-Vac. Proteins were digested by incubating gel pieces with 25 mM Triethylammonium bicarbonate (TEAB) containing 5 μg/mL Trypsin (ThermoFisher #90058), at 30°C, 1000 rpm, overnight. An equivalent volume of 100% ACN was added to enhance digestion and incubated for 15 mins. Supernatants were separated, and gel pieces resuspended in a further 50% ACN/2.5% formic acid for 10 mins, then the supernatants were combined and dried by Speed-Vac.

### p-MS MASCOT

Phosphoproteomic analysis by liquid chromatography mass spectrometry (LC-MS/MS). LC separations were performed with a Thermo Dionex Ultimate 3000 RSLC Nano liquid chromatography instrument using 0.1% formic acid as buffer A and 80% acetonitrile with 0.08% formic acid as buffer B. The peptide samples were loaded on C18 trap columns with 3% acetonitrile / 0.1% trifluoracetic acid at a flow rate of 5 μL/min. Peptide separations were performed over EASY-Spray column (C18, 2 μM, 75 μm x 50 cm) with an integrated nano electrospray emitter at a flow rate of 300 nL/min. Peptides were separated with a 60 min segmented gradient starting from 3%~7% buffer B over 5 min, 7%~25% buffer B over 43 mins, 25%~35% buffer B over 5 min, then raised to 95% buffer B over 3 min and held for 2 min. Eluted peptides were analysed on an Orbitrap Exploris 480 (ThermoFisher Scientific, San Jose, CA) mass spectrometer. Spray voltage was set to 2 kV, RF lens level was set at 40%, and ion transfer tube temperature was set to 275 °C. The mass spectrometer was operated in data-dependent mode (ID low load) with 2 seconds per cycle. The full scan was performed in the range of 350—1200 m/z at nominal resolution of 60,000 at 200 m/z and AGC set to 300% with a custom maximal injection time of 28 ms, followed by selection of the most intense ions above an intensity threshold of 10000 for higher-energy collision dissociation (HCD) fragmentation. HCD normalised collision energy was set to 30%. Data-dependent MS2 scans were acquired for charge states 2 to 6 using an isolation width of 1.2 m/z and a 30 second dynamic exclusion duration. All MS2 scans were recorded with centroid mode using an AGC target set to standard and a maximal fill time of 100 ms.

The .RAW files obtained from the mass spectrometer were processed by Proteome Discoverer v2.4 (ThermoFisher Scientific) using Mascot v 2.6.2 (Matrix Science) as the search engine. A precursor mass tolerance of 10ppm and fragment tolerance of 0.06 was used. An in-house database (MRC_Database_1) was used with trypsin set as the protease which was allowed a maximum of two missed cleavage sites. Oxidation and dioxidation of methionine, phosphorylation of serine, threonine and tyrosine were set as variable modifications, and carbamidomethylation of cysteine was set as a fixed modification. Phospho-site assignment probability was estimated via Mascot and PhosphoRS3.1 (Proteome Discoverer v.1.4-SP1) or ptmRS (Proteome Discoverer v.2.0)..ptmRS was used as a scoring system for the phospho site identification, with a mass tolerance of 0.5 and neutral loss peaks were considered. Phosphorylation site localisation was considered correct only if peptides had a Mascot delta score and ptmRS probability score above 85%.

## Acknowledgements

We thank C. Gonzalez and M. Krahn for the gift of flies. We thank the Dundee imaging facility for support. Stocks obtained from the Bloomington Drosophila Stock Center (NIH P40OD018537) were used in this study. We thank R. Le Borgne for discussion and C. Roubinet for critical reading of the manuscript. Work in JJ’s was supported by grants from Wellcome (100031/Z/12/A) and the BBSRC (BB/V001353/1 and BB/T017546/1).GMF and EH were supported by a Wellcome Trust/Royal Society Sir Henry Dale Fellowship (211209/Z/18/Z). DHM was supported by a Wellcome Trust/Royal Society Sir Henry Dale Fellowship (211193/Z/18/Z) and a Royal Society (RGS\R2\180284) grant.

## REFERENCES

Alexandre, P., Reugels, A. M., Barker, D., Blanc, E. & Clarke, J. D. 2010. Neurons derive from the more apical daughter in asymmetric divisions in the zebrafish neural tube. Nat Neurosci, 13, 673–9.

Atwood, S. X., Chabu, C., Penkert, R. R., Doe, C. Q. & Prehoda, K. E. 2007. Cdc42 acts downstream of Bazooka to regulate neuroblast polarity through Par-6 aPKC. J Cell Sci, 120, 3200–6.

Atwood, S. X. & Prehoda, K. E. 2009. aPKC phosphorylates Miranda to polarize fate determinants during neuroblast asymmetric cell division. Curr Biol, 19, 723–9.

Bellaiche, Y., Radovic, A., Woods, D. F., Hough, C. D., Parmentier, M. L., O’Kane, C. J., Bryant, P. J. & Schweisguth, F. 2001. The Partner of Inscuteable/Discs-large complex is required to establish planar polarity during asymmetric cell division in Drosophila. Cell, 106, 355–66.

Benton, R. & St Johnston, D. 2003. Drosophila PAR-1 and 14-3-3 inhibit Bazooka/PAR-3 to establish complementary cortical domains in polarized cells. Cell, 115, 691–704.

Bodenmiller, B., Campbell, D., Gerrits, B., Lam, H., Jovanovic, M., Picotti, P., Schlapbach, R. & Aebersold, R. 2008. PhosphoPep--a database of protein phosphorylation sites in model organisms. Nat Biotechnol,26, 1339–40.

Buszczak, M., Paterno, S., Lighthouse, D., Bachman, J., Planck, J., Owen, S., Skora, A. D., Nystul, T. G., Ohlstein, B., Allen, A., Wilhelm, J. E., Murphy, T. D., Levis, R. W., Matunis, E., Srivali, N., Hoskins, R. A. & Spradling, A. C. 2007. The carnegie protein trap library: a versatile tool for Drosophila developmental studies. Genetics, 175, 1505–1531.

Cai, Y., Yu, F., Lin, S., Chia, W. & Yang, X. 2003. Apical complex genes control mitotic spindle geometry and relative size of daughter cells in Drosophila neuroblast and pI asymmetric divisions. Cell, 112, 51–62.

Darnat, P., Burg, A., Salle, J., Lacoste, J., Louvet-Vallee, S., Gho, M. & Audibert, A. 2022. Cortical Cyclin A controls spindle orientation during asymmetric cell divisions in Drosophila. Nat Commun, 13, 2723.

Das, R. M. & Storey, K. G. 2012. Mitotic spindle orientation can direct cell fate and bias Notch activity in chick neural tube. EMBO Rep, 13, 448–54.

Deluca, S. Z. & Spradling, A. C. 2018. Efficient Expression of Genes in the Drosophila Germline Using a UAS Promoter Free of Interference by Hsp70 piRNAs. Genetics, 209, 381–387.

Dickinson, D. J., Schwager, F., Pintard, L., Gotta, M. & Goldstein, B. 2017. A Single-Cell Biochemistry Approach Reveals PAR Complex Dynamics during Cell Polarization. Dev Cell, 42, 416–434 e11.

Dumont, N. A., Wang, Y. X., Von Maltzahn, J., Pasut, A., Bentzinger, C. F., Brun, C. E. & Rudnicki, M. A. 2015. Dystrophin expression in muscle stem cells regulates their polarity and asymmetric division. Nat Med, 21, 1455–63.

Emery, G., Hutterer, A., Berdnik, D., Mayer, B., Wirtz-Peitz, F., Gaitan, M. G. & Knoblich, J. A. 2005. Asymmetric Rab 11 endosomes regulate delta recycling and specify cell fate in the Drosophila nervous system. Cell, 122, 763–773.

Fichelson, P., Audibert, A., Simon, F. & Gho, M. 2005. Cell cycle and cell-fate determination in Drosophila neural cell lineages. Trends Genet, 21, 413–20.

Fichelson, P. & Gho, M. 2004. Mother-daughter precursor cell fate transformation after Cdc2 down-regulation in the Drosophila bristle lineage. Dev Biol, 276, 367–77.

Gallaud, E., Pham, T. & Cabernard, C. 2017. Drosophila melanogaster Neuroblasts: A Model for Asymmetric Stem Cell Divisions. Results and problems in cell differentiation, 61, 183–210.

Gillies, T. E. & Cabernard, C. 2011. Cell division orientation in animals. Current biology: CB, 21, R599–609.

Goehring, N. W. 2014. PAR polarity: from complexity to design principles. Experimental cell research, 328, 258–266.

Gogl, G., Tugaeva, K. V., Eberling, P., Kostmann, C., Trave, G. & Sluchanko, N. N. 2021. Hierarchized phosphotarget binding by the seven human 14-3-3 isoforms. Nat Commun, 12, 1677.

Goldstein, B. & Macara, I. G. 2007. The PAR proteins: fundamental players in animal cell polarization. Developmental Cell, 13, 609–622.

Hannaford, M., Loyer, N., Tonelli, F., Zoltner, M. & Januschke, J. 2019. A chemical-genetics approach to study the role of atypical Protein Kinase C in Drosophila. Development, 146, dev170589.

Hannaford, M. R., Ramat, A., Loyer, N. & Januschke, J. 2018. aPKC-mediated displacement and actomyosin-mediated retention polarize Miranda in Drosophila neuroblasts. eLife, 7, 166.

Hayashi, K., Yonemura, S., Matsui, T., Tsukita, S. & Tsukita, S. 1999. Immunofluorescence detection of ezrin/radixin/moesin (ERM) proteins with their carboxyl-terminal threonine phosphorylated in cultured cells and tissues Application of a novel fixation protocol using trichloroacetic acid (TCA) as a fixative. Journal of Cell Science, 112, 1149–1158.

Hilger, M., Bonaldi, T., Gnad, F. & Mann, M. 2009. Systems-wide analysis of a phosphatase knock-down by quantitative proteomics and phosphoproteomics. Mol Cell Proteomics, 8, 1908–20.

Hochegger, H., Takeda, S. & Hunt, T. 2008. Cyclin-dependent kinases and cell-cycle transitions: does one fit all? Nat Rev Mol Cell Biol, 9, 910–6.

Houssin, E., Pinot, M., Bellec, K. & Le Borgne, R. 2021. Par3 cooperates with Sanpodo for the assembly of Notch clusters following asymmetric division of Drosophila sensory organ precursor cells. Elife, 10.

Hu, Y., Sopko, R., Chung, V., Foos, M., Studer, R. A., Landry, S. D., Liu, D., Rabinow, L., Gnad, F., Beltrao, P. & Perrimon, N. 2019. iProteinDB: An Integrative Database of Drosophila Post-translational Modifications. G3 (Bethesda, Md.), 9, 1–11.

Izumi, Y., Ohta, N., Itoh-Furuya, A., Fuse, N. & Matsuzaki, F. 2004. Differential functions of G protein and Baz-aPKC signaling pathways in Drosophila neuroblast asymmetric division. J Cell Biol, 164, 729–38.

Januschke, J. & Loyer, N. 2020. Applications of Immobilization of Drosophila Tissues with Fibrin Clots for Live Imaging. J Vis Exp.

Jones, T. A. & Metzstein, M. M. 2011. A novel function for the PAR complex in subcellular morphogenesis of tracheal terminal cells in Drosophila melanogaster. Genetics, 189, 153–64.

Kemphues, K. 2000. PARsing embryonic polarity. Cell, 101, 345–348.

Khazaei, M. R. & Püschel, A. W. 2009. Phosphorylation of the par polarity complex protein Par3 at serine 962 is mediated by aurora a and regulates its function in neuronal polarity. Journal of Biological Chemistry, 284, 33571–33579.

Kim, A. J. & Griffin, E. E. 2020. PLK-1 Regulation of Asymmetric Cell Division in the Early C. elegans Embryo. Front Cell Dev Biol, 8, 632253.

Kullmann, L. & Krahn, M. P. 2018. Redundant regulation of localization and protein stability of DmPar3. Cellular and molecular life sciences: CMLS, 75, 3269–3282.

Le Borgne, R. & Schweisguth, F. 2003. Unequal segregation of Neuralized biases Notch activation during asymmetric cell division. Dev Cell, 5, 139–48.

Lee, C. Y., Andersen, R. O., Cabernard, C., Manning, L., Tran, K. D., Lanskey, M. J., Bashirullah, A. & Doe, C. Q. 2006. Drosophila Aurora-A kinase inhibits neuroblast self-renewal by regulating aPKC/Numb cortical polarity and spindle orientation. Genes Dev, 20, 3464–74.

Lin, S., Wang, C., Zhou, J., Shi, Y., Ruan, C., Tu, Y., Yao, L., Peng, D. & Xue, Y. 2021. EPSD: a well-annotated data resource of protein phosphorylation sites in eukaryotes. Brief Bioinform, 22, 298–307.

Lopez, M. S., Kliegman, J. I. & Shokat, K. M. 2014. The logic and design of analog-sensitive kinases and their small molecule inhibitors. Methods in enzymology, 548, 189–213.

Lowe, E. D., Tews, I., Cheng, K. Y., Brown, N. R., Gul, S., Noble, M. E. M., Gamblin, S. J. & Johnson, L. N. 2002. Specificity Determinants of Recruitment Peptides Bound to Phospho-CDK2/Cyclin A. Biochemistry, 41, 15625–15634.

Mcgrath, D. A., Balog, E. R., Koivomagi, M., Lucena, R., Mai, M. V., Hirschi, A., Kellogg, D. R., Loog, M. & Rubin, S. M. 2013. Cks confers specificity to phosphorylation-dependent CDK signaling pathways. Nat Struct Mol Biol,20, 1407–14.

Morais-de-Sá, E., Mirouse, V. & St Johnston, D. 2010. aPKC phosphorylation of Bazooka defines the apical/lateral border in Drosophila epithelial cells. Cell, 141, 509–523.

Motegi, F. & Sugimoto, A. 2006. Sequential functioning of the ECT-2 RhoGEF, RHO-1 and CDC-42 establishes cell polarity in Caenorhabditis elegans embryos. Nat Cell Biol, 8, 978–85.

Müller, H. A. & Wieschaus, E. 1996. armadillo, bazooka, and stardust are critical for early stages in formation of the zonula adherens and maintenance of the polarized blastoderm epithelium in Drosophila. Journal of Cell Biology, 134, 149–163.

Munro, E., Nance, J. & Priess, J. R. 2004. Cortical flows powered by asymmetrical contraction transport PAR proteins to establish and maintain anterior-posterior polarity in the early C. elegans embryo. Developmental Cell, 7, 413–424.

Niiya, F., Xie, X., Lee, K. S., Inoue, H. & Miki, T. 2005. Inhibition of cyclin-dependent kinase 1 induces cytokinesis without chromosome segregation in an ECT2 and MgcRacGAP-dependent manner. The Journal of biological chemistry, 280, 36502–36509.

Noatynska, A., Tavernier, N., Gotta, M. & Pintard, L. 2013. Coordinating cell polarity and cell cycle progression: what can we learn from flies and worms? Open Biol, 3, 130083.

Nurse, P. 1997. Regulation of the eukaryotic cell cycle. Eur J Cancer, 33, 1002–4.

Oon, C. H. & Prehoda, K. E. 2019. Asymmetric recruitment and actin-dependent cortical flows drive the neuroblast polarity cycle. eLife, 8, 723.

Petronczki, M. & Knoblich, J. A. 2001. DmPAR-6 directs epithelial polarity and asymmetric cell division of neuroblasts in Drosophila. Nature Cell Biology, 3, 43–49.

Potapova, T. A., Daum, J. R., Pittman, B. D., Hudson, J. R., Jones, T. N., Satinover, D. L., Stukenberg, P. T. & Gorbsky, G. J. 2006. The reversibility of mitotic exit in vertebrate cells. Nature, 440, 954–8.

Potapova, T. A., Sivakumar, S., Flynn, J. N., Li, R. & Gorbsky, G. J. 2011. Mitotic progression becomes irreversible in prometaphase and collapses when Wee1 and Cdc25 are inhibited. Mol Biol Cell, 22, 1191–206.

Ramat, A., Hannaford, M. & Januschke, J. 2017. Maintenance of Miranda Localization in Drosophila Neuroblasts Involves Interaction with the Cognate mRNA. Current biology: CB, 27, 2101–2111.e5.

Reich, J. D., Hubatsch, L., Illukkumbura, R., Peglion, F., Bland, T., Hirani, N. & Goehring, N. W. 2019. Regulated Activation of the PAR Polarity Network Ensures a Timely and Specific Response to Spatial Cues. Curr Biol, 29, 1911–1923 e5.

Rodriguez, J., Peglion, F., Martin, J., Hubatsch, L., Reich, J., Hirani, N., Gubieda, A. G., Roffey, J., Fernandes, A. R., St Johnston, D., Ahringer, J. & Goehring, N. W. 2017. aPKC Cycles between Functionally Distinct PAR Protein Assemblies to Drive Cell Polarity. Developmental Cell, 42, 400–415.e9.

Rolls, M. M., Albertson, R., Shih, H.-P., Lee, C.-Y. & Doe, C. Q. 2003. Drosophila aPKC regulates cell polarity and cell proliferation in neuroblasts and epithelia.Journal of Cell Biology, 163, 1089–1098.

Rose, L. S. & Kemphues, K. J. 1998. Early patterning of the C. elegans embryo. Annu Rev Genet, 32, 521–45.

Schober, M., Schaefer, M. & Knoblich, J. A. 1999. Bazooka recruits Inscuteable to orient asymmetric cell divisions in Drosophila neuroblasts. Nature, 402, 548–551.

Schumacher, J. M., Ashcroft, N., Donovan, P. J. & Golden, A. 1998. A highly conserved centrosomal kinase, AIR-1, is required for accurate cell cycle progression and segregation of developmental factors in Caenorhabditis elegans embryos. Development, 125, 4391–402.

Schweisguth, F. 2015. Asymmetric cell division in the Drosophila bristle lineage: from the polarization of sensory organ precursor cells to Notch-mediated binary fate decision. Wiley interdisciplinary reviews. Developmental biology, 4, 299–309.

Simpson, P. 1990. Lateral inhibition and the development of the sensory bristles of the adult peripheral nervous system of Drosophila. Development, 109, 509–19.

Songyang, Z., Blechner, S., Hoagland, N., Hoekstra, M. F., Piwnica-Worms, H. & Cantley, L. C. 1994. Use of an oriented peptide library to determine the optimal substrates of protein kinases. Curr Biol, 4, 973–82.

Sunchu, B. & Cabernard, C. 2020. Principles and mechanisms of asymmetric cell division. Development, 147.

Suzuki, K., Sako, K., Akiyama, K., Isoda, M., Senoo, C., Nakajo, N. & Sagata, N. 2015. Identification of non-Ser/Thr-Pro consensus motifs for Cdk1 and their roles in mitotic regulation of C2H2 zinc finger proteins and Ect2. Sci Rep, 5, 7929.

Tio, M., Udolph, G., Yang, X. & Chia, W. 2001. cdc2 links the Drosophila cell cycle and asymmetric division machineries. Nature, 409, 1063–1067.

Usui, K. & Kimura, K.-I. 1992. Sensory mother cells are selected from among mitotically quiescent cluster of cells in the wing disc of Drosophila. Development, 116, 601–610.

Van Horn, R. D., Chu, S., Fan, L., Yin, T., Du, J., Beckmann, R., Mader, M., Zhu, G., Toth, J., Blanchard, K. & Ye, X. S. 2010. Cdk1 Activity Is Required for Mitotic Activation of Aurora A during G2/M Transition of Human Cells. Journal of Biological Chemistry, 285, 21849–21857.

Wang, H., Ouyang, Y., Somers, W. G., Chia, W. & Lu, B. 2007. Polo inhibits progenitor self-renewal and regulates Numb asymmetry by phosphorylating Pon. Nature, 449, 96–100.

Wang, H., Somers, G. W., Bashirullah, A., Heberlein, U., Yu, F. & Chia, W. 2006. Aurora-A acts as a tumor suppressor and regulates self-renewal of Drosophila neuroblasts. Genes & Development, 20, 3453–3463.

Wirtz-Peitz, F., Nishimura, T. & Knoblich, J. A. 2008. Linking cell cycle to asymmetric division: Aurora-A phosphorylates the Par complex to regulate Numb localization. Cell, 135, 161–173.

Wodarz, A., Ramrath, A., Grimm, A. & Knust, E. 2000. Drosophila atypical protein kinase C associates with Bazooka and controls polarity of epithelia and neuroblasts. Journal of Cell Biology, 150, 1361–1374.

Wodarz, A., Ramrath, A., Kuchinke, U. & Knust, E. 1999. Bazooka provides an apical cue for Inscuteable localization in Drosophila neuroblasts. Nature, 402, 544–547.

Wu, J., Tannan, N. B., Vuong, L. T., Koca, Y., Collu, G. M. & Mlodzik, M. 2023a. Par3/Bazooka binds NICD and promotes Notch signalling during <EM>Drosophila</EM> development. bioRxiv, 2022.05.24.493322.

Wu, S., Yang, Y., Tang, R., Zhang, S., Qin, P., Lin, R., Rafel, N., Lucchetta, E. M., Ohlstein, B. & Guo, Z. 2023b. Apical-basal polarity precisely determines intestinal stem cell number by regulating Prospero threshold. Cell Rep, 42, 112093.

Yu, F., Morin, X., Cai, Y., Yang, X. & Chia, W. 2000. Analysis of partner of inscuteable, a novel player of Drosophila asymmetric divisions, reveals two distinct steps in inscuteable apical localization. Cell, 100, 399–409.

Zhai, B., Villen, J., Beausoleil, S. A., Mintseris, J. & Gygi, S. P. 2008. Phosphoproteome analysis of Drosophila melanogaster embryos. J Proteome Res, 7, 1675–82.

