## Supplementary figures and images for "Asymmetric cell division-specific phosphorylation of PAR-3 regulates neuroblasts polarisation and sensory organ formation in *Drosophila*"

### Fig 2 supplement

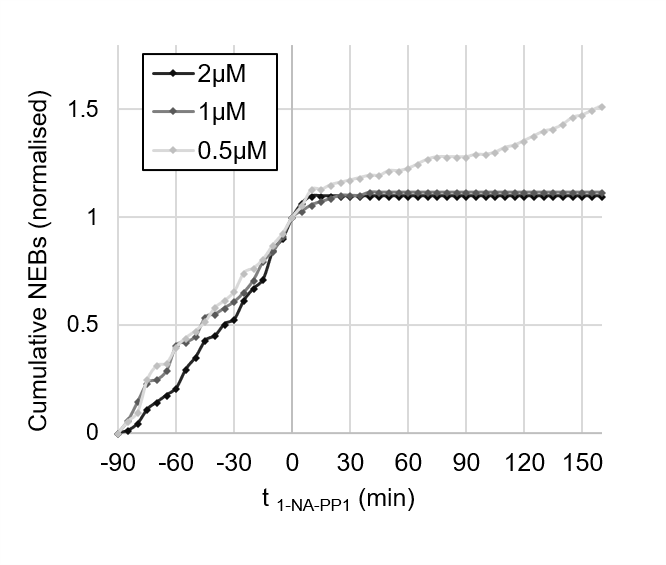

### Fig 3 supplement

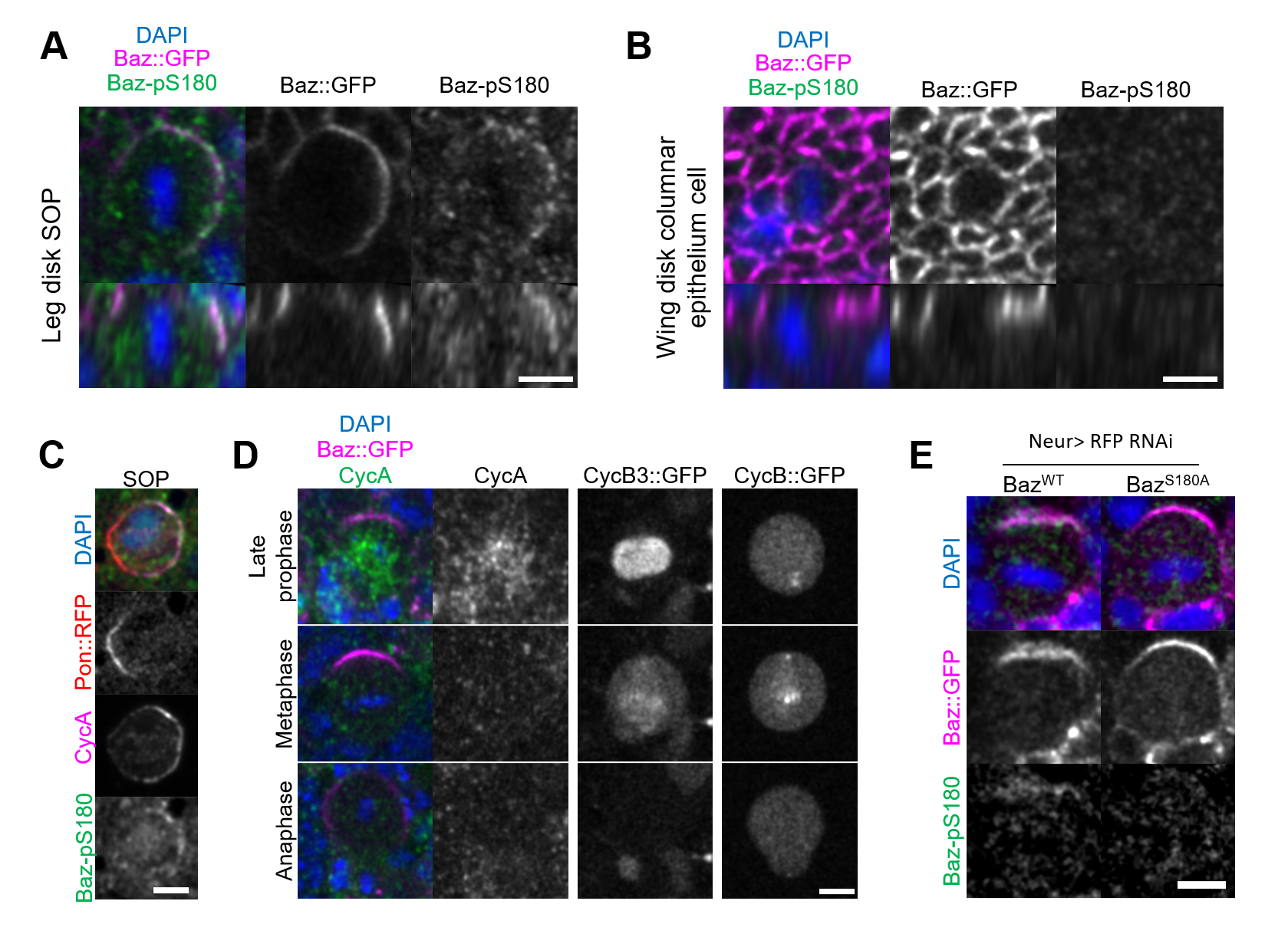
